# An asymmetric sheath controls flagellar supercoiling and motility in the Leptospira spirochete

**DOI:** 10.1101/847533

**Authors:** Kimberley H. Gibson, Felipe Trajtenberg, Elsio A. Wunder, Megan R. Brady, Fabiana San Martin, Ariel E. Mechaly, Zhiguo Shang, Jun Liu, Mathieu Picardeau, Albert I. Ko, Alejandro Buschiazzo, Charles V. Sindelar

## Abstract

Spirochete bacteria, including important pathogens, exhibit a distinctive means of swimming via undulations of the entire cell. Motility is powered by the rotation of supercoiled ‘endoflagella’ that wrap around the cell body, confined within the periplasmic space. To investigate the structural basis of flagellar supercoiling, which is critical for motility, we determined the structure of native flagellar filaments from the spirochete *Leptospira* by integrating high-resolution cryo-electron tomography and X-ray crystallography. We show that these filaments are coated by a highly asymmetric, multi-component sheath layer, contrasting with flagellin-only homopolymers previously observed in exoflagellated bacteria. Distinct sheath proteins localize to the filament inner and outer curvatures to define the supercoiling geometry, explaining a key functional attribute of the spirochete flagellum.

**One Sentence summary:** The corkscrew-like motility of Spirochete bacteria is enabled by a unique, asymmetrically constructed flagellum that wraps around the cell body within the periplasm.

## Main Text

The spirochetes, double-membrane bacteria with helically coiled cells, are a diverse phylum that includes the agents of leptospirosis (*Leptospira* spp.), syphilis (*Treponema pallidum*) and Lyme disease (*Borrelia burgdorferi*). Spirochetes have powerful swimming capabilities that enable them to rapidly disseminate through connective tissue, blood, and organs(Wunder, Figueira, Santos, et al., 2016). This swimming capability is enabled by a unique configuration of the flagellum in these organisms(Charon et al., 2012; Li, Corum, et al., 2000; Wolgemuth, Charon, Goldstein, & Goldstein, 2006) (Fig. 1A), which allows them to ‘drill’ through highly viscous media(Li, Motaleb, Sal, Goldstein, & Charon, 2000; Wolgemuth et al., 2006). By housing their flagella in the periplasm, spirochetes protect these filaments from environmental insults, such as immune surveillance(Vernel-Pauillac & Werts, 2018). All known spirochetal flagella exhibit supercoiled architectures(Charon et al., 1992; Dombrowski et al., 2009), which have been linked to productive motility(Wolgemuth, 2015), and amongst pathogenic species, to virulence(Ko, Goarant, & Picardeau, 2009; McBride, Athanazio, Reis, & Ko, 2005; Sultan et al., 2013; Wunder, Figueira, Benaroudj, et al., 2016).

**Figure 1.**
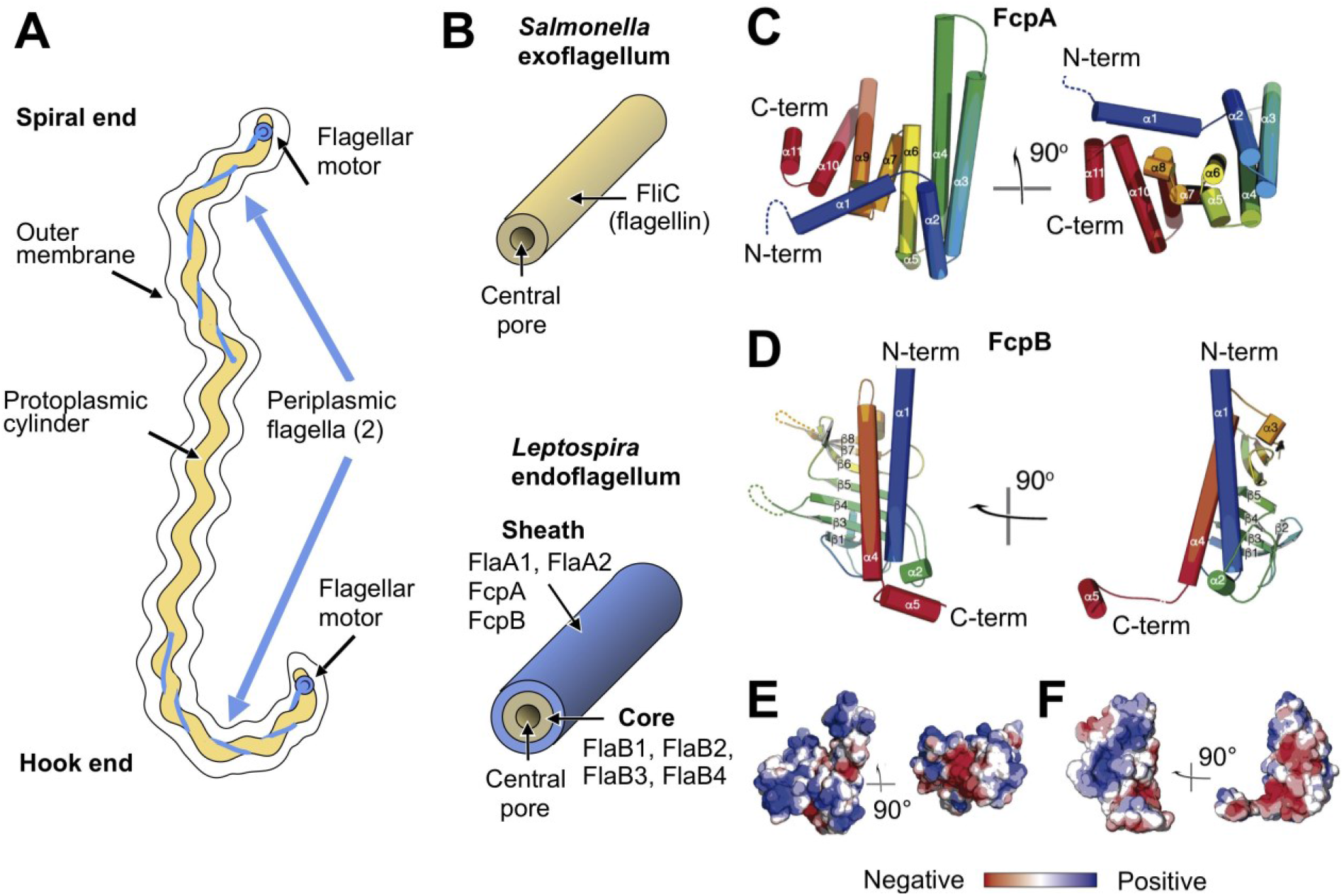
X-ray crystal structures of the sheath proteins, FcpA from *L. biflexa* and FcpB from *L. interrogans*. **A**. Schematic of a *Leptospira* cell. Each cell has two flagella sandwiched between the inner and outer membranes; a single flagellum extends from a motor at either end. **B**. Comparison of the predicted endoflagellar filament composition and morphology in *Leptospira* with the *Salmonella* exoflagellum. **C.** Structure of *L. biflexa* FcpA in two orthogonal views, depicted as a cartoon colored with a ramp from blue (N-terminus) to red (C-terminus). The dotted line stands for the flexible 54 amino acids at the N-terminus, not visible in electron density. **D.** Structure of *L. interrogans* FcpB in two orthogonal views, similar color code as panel A. **E.** Solvent accessible surface of FcpA colored according to an electrostatic potential ramp from negative (red) through neutral (white) to positive (blue) potentials. **F.** Solvent accessible surface of FcpB, similar color code as panel E. The perspective was chosen to show the interacting surface of FcpB (positively charged region, panel F left-hand side), in an open-book view (approximately 180º rotated according to a vertical axis), with FcpA (negative cleft, panel E right-hand side). This interaction was later uncovered by studying the whole filament assembly (see Fig. 4E).

*Leptospira*, which includes pathogenic species that cause human and animal disease and saprophytic species(Picardeau, 2017), is unique among bacteria, including spirochetes, in that isolated flagellar filaments spontaneously curl resembling lassos, following tightly supercoiled, nearly coplanar paths(Bromley & Charon, 1979) (Supplementary Fig. 1). The flagellar filament thus superposes a curved path on the underlying helical body, resulting in overall ‘spiral’ or ‘hook’ cell end shapes depending on the motor rotation direction(Charon & Goldstein, 2002) (Fig. 1A). Similar to other spirochetes, *Leptospira* travel rapidly and unidirectionally when flagellar motors at opposing ends of the organism rotate in opposite directions (i.e. counterclockwise *vs.* clockwise when viewing the motor from the cell exterior), but not when the motors are all rotating in the same direction(Goldstein & Charon, 1988; Wolgemuth, 2015). *Leptospira* can also adopt a distinct ‘crawling’ mode while bound to surfaces(Tahara et al., 2018).

As with all spirochetes, *Leptospira* have a complex flagellar composition compared to exoflagellated bacteria(Charon & Goldstein, 2002) (Fig. 1B, Supplementary Table 1, Supplementary Fig. 2). Flagellar filaments from non-spirochete bacteria generally have a single protein component(Erhardt, Namba, & Hughes, 2010) (flagellin). In contrast, spirochetal flagellar filaments are comprised of a flagellin homolog (FlaB) and FlaA(Brahamsha & Greenberg, 1988), a second and completely unrelated component. FlaA and FlaB proteins can occur in multiple isoforms within a single organism(Wolgemuth et al., 2006) (Supplementary Table 1). Deletion of FlaA or FlaB in *Leptospira* leads to distinct phenotypes: deletion of FlaA affects flagellar curvature and diminishes motility and pathogenicity(Lambert et al., 2012), while deletion of FlaB eliminates the filament entirely(Picardeau, Brenot, & Saint Girons, 2001). Similar observations have been made in other spirochetes(Li, Corum, et al., 2000; Motaleb et al., 2000), indicating that the structural features of *Leptospira* flagella represent important functional adaptations relevant to spirochetes as a whole. However, *Leptospira* flagella have unique features not found in other spirochetes. Recently, two novel components of the *Leptospira* flagellum, FcpA and FcpB (Supplementary Table 1), were identified whose deletion abolishes the ability of flagella to assume their characteristic supercoiled form and at the same time dramatically reduces motility and virulence(Wunder, Figueira, Benaroudj, et al., 2016; Wunder et al., 2018).

The detailed structural underpinnings of the endoflagellar filament and its components, has remained mysterious for *Leptospira* or indeed, for spirochetes in general. Here, we present structures of flagellar filaments from the spirochete *Leptospira biflexa*, a saprophytic species, solved by a combination of cryo-tomography, sub-tomogram averaging, X-ray crystallography and molecular docking. Our structures reveal that a conserved FlaB core assembly is enclosed by an asymmetric assembly of sheath proteins with novel folds and functions. Analysis of mutant filaments, which have lost one or more of these sheath elements, indicates that the sheath enforces different core lattice geometries on different sides of the filament, thus promoting a supercoiled flagellar shape. This asymmetric supercoiling mechanism may be relevant to other spirochetes, contributing to their distinctive swimming modes.

## FcpA and FcpB X-ray structures

Using X-ray crystallography, we established that the flagellar sheath components FcpA and FcpB adopt unique folds, unrelated to the globular domains found in typical flagellins (Fig. 1C-F). FcpA from *L. biflexa* was crystallized in three different space groups(San Martin et al., 2017) (Supplementary Table 2), revealing a 3D fold previously unseen within the PDB (release July 2019) as reported by several structural alignment algorithms(Holm & Laakso, 2016; Madej et al., 2014). FcpB from the pathogenic species, *L. interrogans* (ortholog of the *L. biflexa* protein; Supplementary Table 1, Supplementary Fig. 2), was crystallized in the orthorhombic space group P2_1_2_1_2_1_ (Supplementary Table 2). The resulting fold exhibits distant structural homology to proteins of the Mam33 family, including human mitochondrial protein p32(Jiang, Zhang, Krainer, & Xu, 1999) that exhibit affinity for a number of protein partners. Homology to FcpB was also detected with VC1805(Sheikh et al., 2008), a protein from *Vibrio cholerae* that can bind complement protein C1q.

FcpA and FcpB are highly distinct in overall size and shape. FcpA assumes an all-helical fold consisting of a twisted stack of eleven α-helices, arranged in a ‘V’-shaped architecture with arms of unequal length (Fig. 1C). The long arm is formed by an α-helical hairpin structure (α3-4) emanating from a 4-helix bundle (comprising helices α2-3-4-6), while the shorter arm is mainly composed of two 3-helix bundles (α6-7-9 and α9-10-11) bridging to the longer arm via a helical hairpin (α9-10). In contrast to FcpA, FcpB adopts a wide and flat oblong shape, consisting of an 8-stranded anti-parallel β-sheet with two long α-helices (α1 and α4) packed on one side (Fig. 1D). These major structural differences between FcpA and FcpB are evident even at relatively low resolution (15-20Å), facilitating their identification in the tomographic reconstructions described below.

## 3D reconstruction of flagellar filaments

The lasso-like supercoiling geometry of intact *Leptospira* flagellar filaments precluded the use of most structure determination methods, including cryo-EM single-particle 3D reconstruction (Supplementary Fig. 1, Methods), motivating us to apply a cryo-electron tomography approach. Overlapping series of cubic sub-volumes following curved filament paths were manually selected from 62 reconstructed tomographic volumes from samples of purified, wild-type *L. biflexa* flagellar filaments. A total of ~10,800 sub-volumes were input to a customized subtomogram alignment procedure (Supplementary Figs. 3, 4, Methods). The resulting averaged map achieved an overall (average) resolution of ~10Å (Supplementary Fig. 5) and exhibits a series of concentric protein layers centered around a ~2 nm central pore, as well as a distinct array of globular features decorating the filament surface (Fig. 2). The morphology of the inner two density layers (Fig. 2F) closely resembles that of helically assembled flagellin domains D0 and D1 of filaments from exoflagellated bacteria(Wang et al., 2017; Yonekura, Maki-Yonekura, & Namba, 2003). This structural homology is consistent with earlier predictions that the core of the spirochete filament is composed of FlaB(Li, Wolgemuth, Marko, Morgan, & Charon, 2008; Nauman, Holt, & Cox, 1969), which is homologous to flagellin but lacks the D2 and D3 domains (Supplementary Fig. 6).

**Figure 2.**
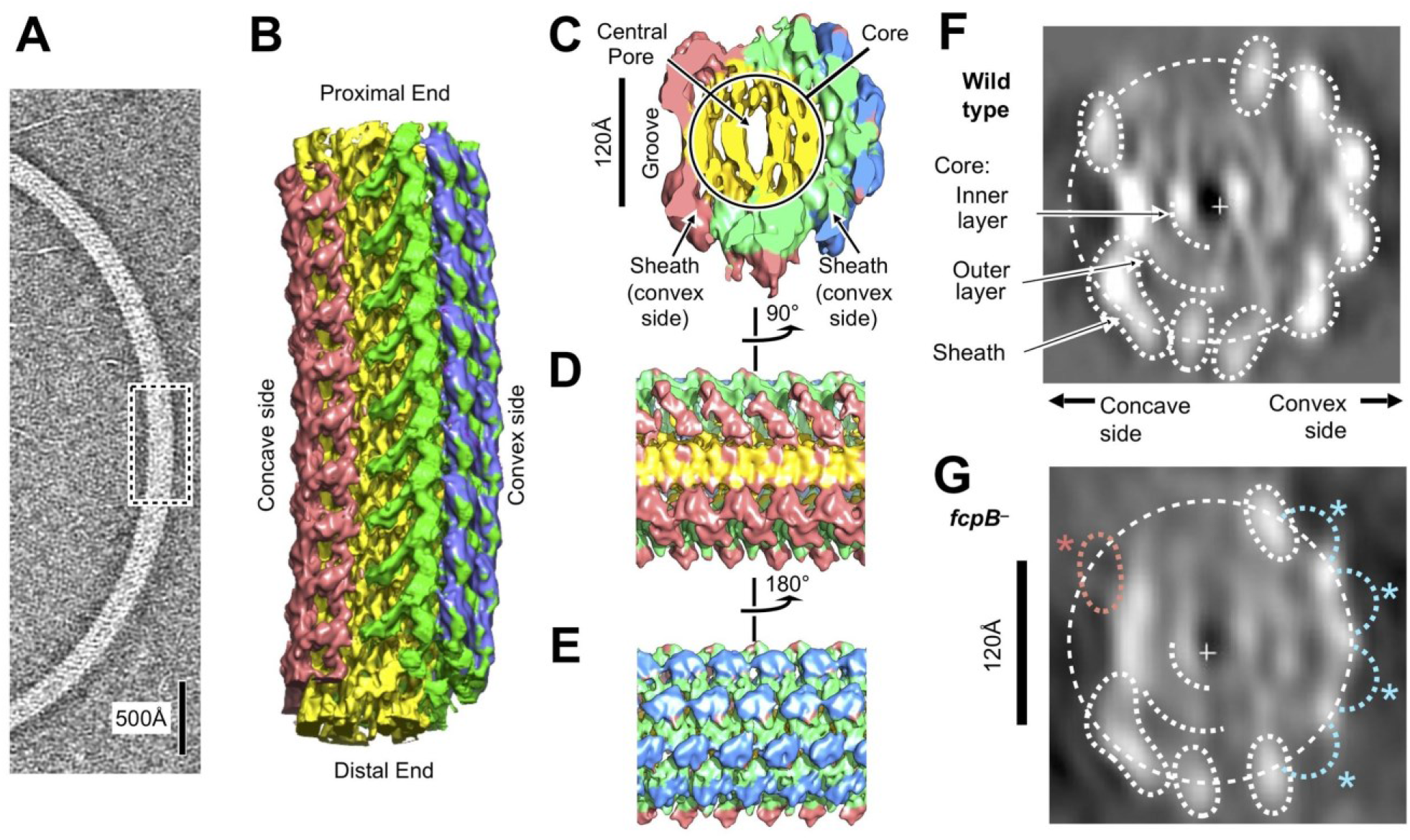
Flagellar filament purified from wild-type *L. biflexa* Patoc with asymmetric sheath layer visualized by cryo-tomographic sub-tomogram averaging. **A.** Cryo-tomographic slice of the flagellar filament in vitrified ice. Dashed box denotes the approximate dimensions extracted for sub-tomographic averaging. **B.** Final averaged volume (isosurface representation) of the flagellar filament denoting segmented regions of sheath (red, green, and blue) and core (yellow). The red sheath regions are located on the inner curvature or ‘concave’ side of the filament, whereas the blue and green sheath regions are located on the outer curvature, or ‘convex’ side. **C.** Cross-sectional view of the sub-tomographic average; the filament diameter ranges from 210-230Å. **D-E.** Rotated views of the sub-tomographic average. **F**, Projected wild-type map cross-section, filtered to 18Å resolution, showing features corresponding to core and sheath elements. **G.** Projected *fcpB*^−^ map cross-section, highlighting differences with the wild-type projection in **F.** Four missing densities on the convex side (blue asterisks) correspond to fitted locations of FcpB in the wild-type map; an additional missing density on the concave side (red asterisk) is provisionally assigned as FlaA1 and/or FlaA2 (see text).

## Asymmetric sheath composition

The tomographic map identified sheath density features that surrounded the two core layers but were not uniformly distributed. Sheath densities were mainly localized on the outer curvature of the filament, or ‘convex’ side, thus offsetting the filament center of mass away from the central pore when the filament is viewed in cross section (Fig. 2C, F, Supplementary Video 1). The finding of a break in the helical symmetry in the *Leptospira* endoflagella sharply contrasts with symmetric structures which were previously observed for exoflagella from *Salmonella*(Sasaki et al., 2018; Wang et al., 2017; Yonekura et al., 2003), *Campylobacter*(Galkin et al., 2008), *Pseudomonas aeruginosa* and *Bacillus subtilis*(Wang et al., 2017).

We used two independent docking strategies to fit FcpA and FcpB molecular envelopes into the reconstructed filament map (see Methods) and identified convergent locations for these two proteins that were distributed across the convex side of the sheath region (Fig. 3, Supplementary Video 1). These sheath sites form an array of globular features that closely follows the canonical 11-protofilament helical lattice identified in flagellar filaments of *Salmonella* and other non-spirochete bacteria (Supplementary Fig. 7). For FcpA, six adjacent, curvilinear rows of ‘V’- shaped density profiles form an interlocking array (Fig. 3, Supplementary Fig. 8). The FcpB sites were resolved as four adjacent rows of flattened density lobes protruding slightly outward from the FcpA portion of the sheath. The resulting assembly consists of an inner layer of FcpA molecules in close contact with the FlaB core, and an outer layer of FcpB molecules nestled between rows of FcpA (Supplementary Video 2).

**Figure 3.**
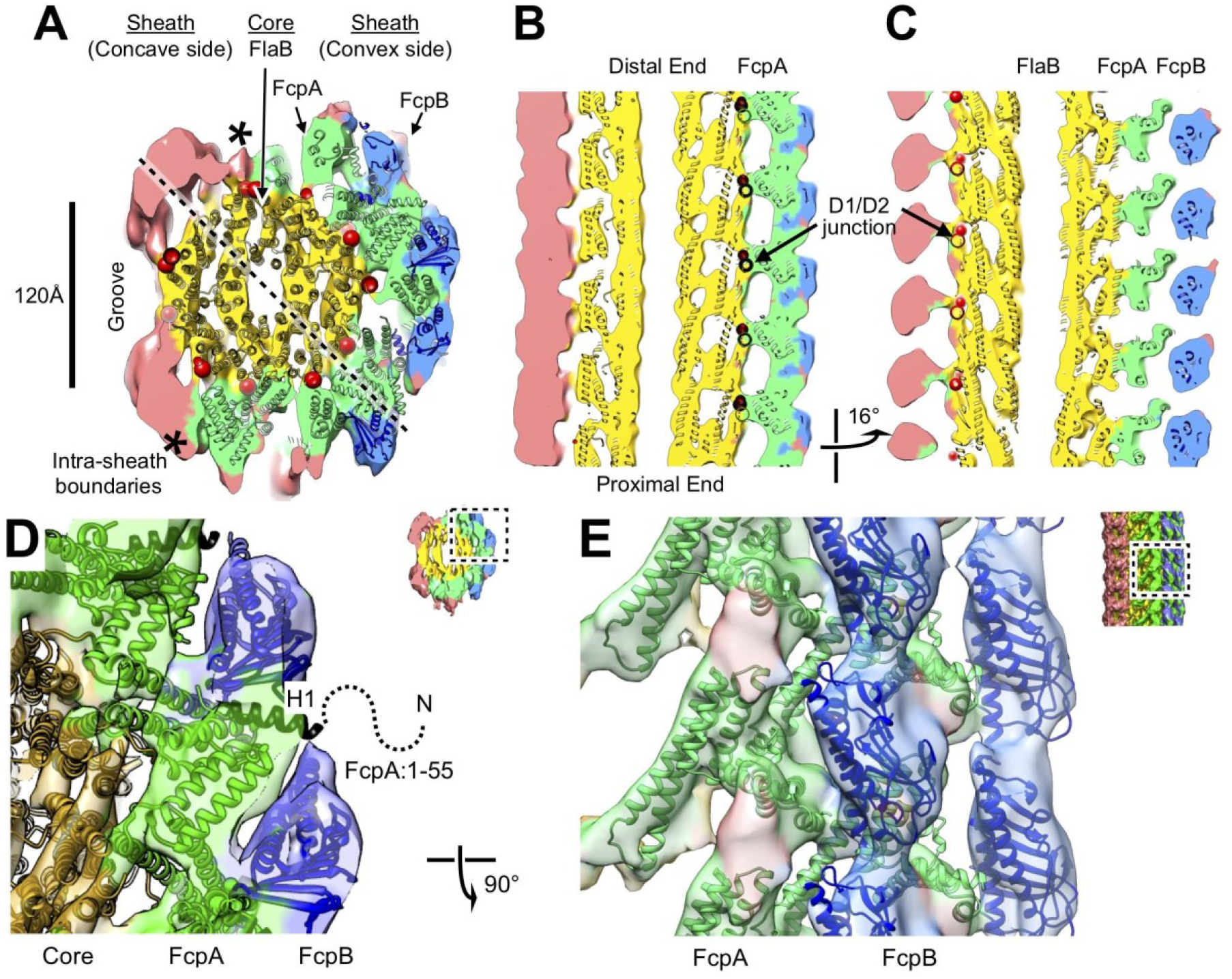
Atomic model of the core and sheath regions of the *L. biflexa* flagellar filament obtained by docking X-ray crystal structures into the cryo-EM map. **A.** Cross-sectional slice of the filament density map isosurface with fitted models of the pseudo-symmetric core assembly (yellow ribbons) and two sheath components, FcpA and FcpB, which localize to the filament outer curvature. Six FcpA protofilaments (green ribbons) directly contact the core and support an outer layer consisting of four FcpB protofilaments (blue ribbons). Asterisks denote boundaries between FcpA and inner curvature density (red). Red markers denote the location of the junction between the modeled D1 α-helical domain of FlaB and the species-specific insertion that substitutes for the D2/D3 outer domains found in *Salmonella* spp. FliC (flagellin) but not in *Leptospira* spp. FlaB. **B.** Longitudinal slice through the filament center, corresponding to the dashed line in A, showing the central channel surrounded by the core and sheath layers. The major interface region between FcpA and the core coincides with this insertion. **C.** A 16° rotated view of the map in **B** showing core-sheath contacts at the site of the FcpB insertions on the opposite side of the filament (concave side); identity of the sheath protein (red) is unassigned. **D.** Close-up cross-sectional view of the averaged filament map showing X-ray model fits of FcpA and FcpB in the outer curvature sheath region. **E**, Close-up view of the averaged filament map rotated 90° relative to the view in **D** showing X-ray model fits.

The asymmetric localization of FcpA and FcpB, specifically on the convex side of the sheath, is distinct from previously characterized exoflagella but is consistent with prior flagellum electron-microscopic imaging studies performed with antibody labels(Wunder, Figueira, Benaroudj, et al., 2016; Wunder et al., 2018). The localization of FcpB sites were further validated by comparison with a similar reconstruction of filaments from an *fcpB*^−^ mutant (Fig. 2G) for which, the projected cross-section of the *fcpB*^−^ map was missing four globular features found in the wild-type map (Fig. 2F) that coincide with our FcpB docking solutions. The signal-to-noise ratio was too low to perform similar sub-tomogram averaging for filaments from an *fcpA*^−^ mutant. However, the filaments from the *fcpA*^−^ mutant were significantly smaller in diameter than filaments from wild-type and *fcpB*^−^ mutants (Supplementary Fig. 9), indicating that filaments from the *fcpA*^−^ mutant have lost FcpA and FcpB(Wunder, Figueira, Benaroudj, et al., 2016).

Additional sheath features are found on the filament concave side, highly divergent from the ones observed on the convex face. These inner-side elements manifested as two rows of elongated density units, separated by a groove that exposes the core (Figs. 2C, 3A). Further modelling of the sheath was precluded as atomic models are not available for FlaA1 and FlaA2, the only other known components of the *Leptospira* flagellar filament. FlaA is also presumed to reside in the spirochete sheath(Li et al., 2008), and comparing the *fcpB*^−^ map with the wild-type map, a missing mass within the concave sheath region indeed suggests candidate positions for FlaA1 and FlaA2 (Fig. 2G, red asterisk). A comprehensive accounting of these features within the concave sheath region awaits further structural and biochemical data.

## Filament core structure

Additional averaging of the map core region revealed homology with flagellar filaments from *Salmonella* and other exoflagellates(Wang et al., 2017). Due to a strongly preferred filament orientation in the specimen ice layer (Supplementary Fig. 1), the potential effect of a ‘missing wedge’ of Fourier-space data characteristic of the cryo-tomography(Carazo, Herman, Sorzano, & Marabini, 2006) could not be fully eliminated. The resulting map resolution anisotropy (Supplementary Fig. 5) interfered with structural analysis of the core region, where globular features were neither evident nor expected based on available models of the flagellar core(Yonekura et al., 2003). By performing 3D auto-correlation analysis within the core region, however, we detected a signal corresponding to the 11-fold helical symmetry operator present in previously determined flagellar structures (Supplementary Fig. 10), congruent with the observed lattice positions of FcpA and FcpB molecules identified in the sheath (Supplementary Fig. 7). Despite minor deviations from this symmetry due to filament curvature, 11-fold averaging using the symmetry operator enabled additional features in the core region to be resolved, including some α-helices, and revealed a structural homology with the D0/D1 region of the *Salmonella* flagellum (Fig. 3B, C, Supplementary Fig. 11, Supplementary Video 2).

Following 11-fold averaging, improved resolution in core region allowed for straightforward fitting of a core atomic model representing a single 52Å repeat of FlaB (11 subunits), which we obtained by applying estimated pseudo-helical symmetry parameters to a FlaB monomer homology model (backbone-only atoms) derived from the reported flagellin atomic structure(Wang et al., 2017; Yamashita et al., 1998; Yonekura et al., 2003) (see Methods). The resulting model also shows good agreement with the non-symmetrized map when the superposition is viewed from favorable orientations that minimize missing wedge artefacts (Fig. 3B, C, Supplementary Video 3).

## Conserved protein-protein interfaces

The resulting filament atomic model exhibits a consistent set of interactions between FcpA and the FlaB core, reflecting the approximate helical symmetry of fitted components (Fig. 4A-C). FcpA molecules project both arms toward the core region, positioning the long arm within contact radius of two predicted loop regions in FlaB (Fig. 4D). Such interactions bury the same protein surface of the FcpA α3-α4 hairpin that forms crystal lattice contacts in all three FcpA X-ray structures. The FlaB loop regions engaged in these interactions project outwards from the core surface toward the predicted FcpA arm positions. Thus, FlaB loops in *Leptospira* (both 6-7 residues in length) likely contribute to the functional interface between FcpA and the core.

**Figure 4.**
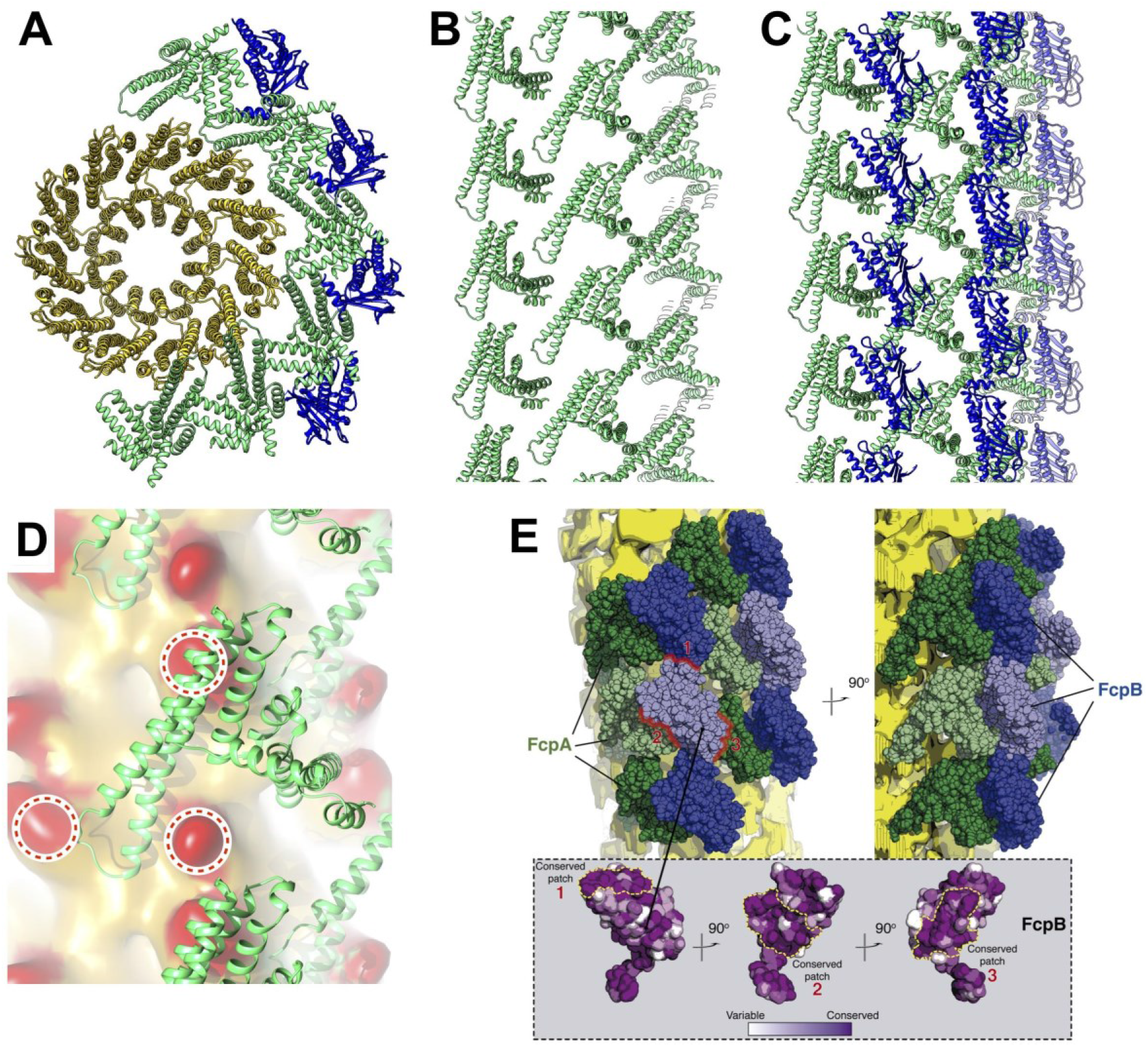
Predicted protein-protein interactions from the combined lattice and sheath model. **A – C.**Ribbon model depiction of the core-sheath atomic model; FlaB is colored gold, FlaA green, FlaB blue. (**A**) is an axial cross-section view, (**B**) is a lateral view of the FcpA lattice only, (**C**) shows the full FcpA/FcpB lattice. **D.** Predicted interactions between FcpA (green ribbon) and predicted loop regions from FlaB (red, circled). **E.** Conserved sequence elements on the FcpB surface are oriented towards neighboring FcpA molecules.

Modeled FcpB subunits project their central β-sheets radially away from the core and present several elements within contact radius of the FcpA inner sheath layer, including one of the long *α*-helices (*α*4) as well as several loops located at the proximal edge of the β-sheet (Fig. 3D). We identified two highly conserved surface patches on FcpB which are in close proximity with nearby FcpA molecules (Fig. 4E), consistent with their likely role as contact interfaces. The corresponding FcpA surface interaction regions predicted by our model are also conserved, and largely match a conserved protein:protein interface found in all FcpA crystal forms. Complementary electrostatic potentials also map onto the FcpA:FcpB interface (Fig. 1E, F). The modelled FcpB sites are too far away from the core to make direct contact, providing an explanation for why deletion of FcpA leads to concurrent loss of FcpB from the filament(Wunder et al., 2018).

The wild-type filament map provides evidence of contact between FcpA and unassigned elements within the concave sheath region, particularly on one side of the groove (Fig 3A; bottom left corner). Since the unassigned material in the concave sheath region is likely composed of FlaA1 and/or FlaA2, FcpA may interact directly with one or both FlaA isoforms. Indeed, pull-down assays of purified *L. biflexa* flagellar filaments recently identified interactions between FcpA and FlaA2(Sasaki et al., 2018).

## The sheath promotes filament curvature

Trajectory analysis of aligned filament subtomograms from wild-type, *fcpA*^−^ and *fcpB*^−^ samples reveals that *fcpB* and *fcpA* deletions cause isolated *Leptospira* flagellar filaments to progressively straighten (Fig. 5A-C). This observation supports and extends previous reports associating these mutations with loss of filament curvature together with pronounced functional effects(Wunder, Figueira, Benaroudj, et al., 2016; Wunder et al., 2018). Wild-type filament segments exhibit a single peak in the measured curvature distribution (Fig. 5A), with an average value of ~5 μm^−1^ (200 nm radius of curvature). In contrast, the average curvature of *fcpA*^−^ filaments (which lack both FcpA and FcpB^4^) is reduced by 40% (~3 μm^−1^, corresponding to 330 nm radius of curvature; Fig. 5C). Mutant *fcpB*^−^ filaments (which still contain FcpA^4^) show a mixed population of curvatures, with approximately half falling within a peak near the wild-type curvature, while the curvature distribution of the other half resembles that of *fcpA*^−^ filaments (Fig. 5B).

**Figure 5.**
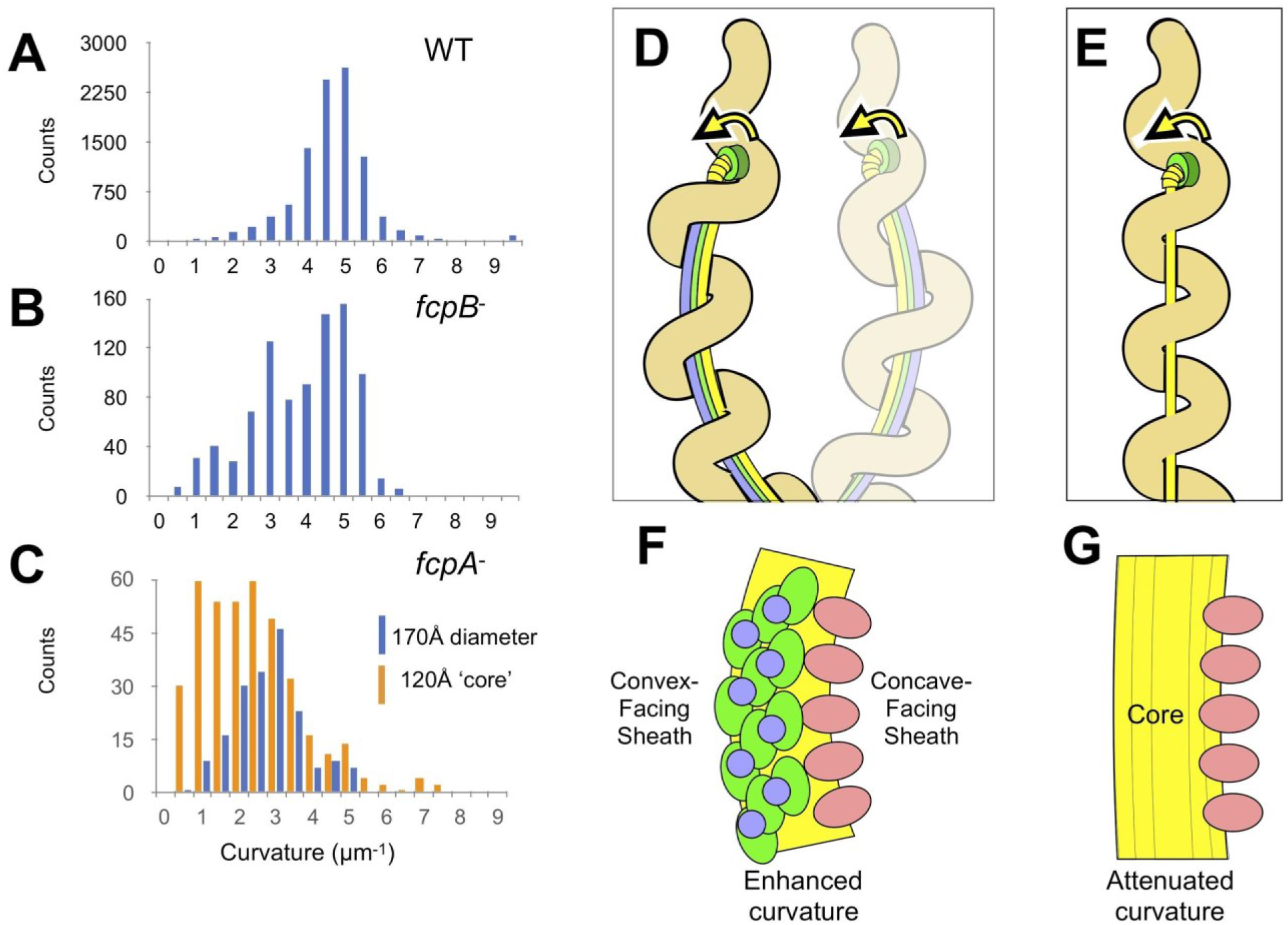
The sheath amplifies flagellar curvature to enable motility in the spirochete *Leptospira* spp. **A**. Histogram of wild-type filament curvatures derived from 3D filament trajectories. A minority of filaments presumed to have shed some or all of the sheath (see Supplementary Fig. 9), as judged by a smaller measured diameter, were excluded from this analysis. **B**. Histogram of *fcpB*^−^ mutant filament curvatures. As in A, a minority population of smaller-diameter filaments were excluded. **C**. Histogram of *fcpA*^−^ mutant filament curvatures, subdivided into distributions for the larger-diameter population (see Supplementary Fig. 9B, 3rd panel) and the smaller-diameter population (see Supplementary Fig. 9B, 4th panel). **D**. Model depicting how sheath-enforced flagellar curvature would interact with the coiled body in *Leptospira* to generate large-scale curved deformations in the body. **E**. If flagellar curvature is lost, the flagellum can pass straight through the body helix without deforming it, so filament rotation does not directly induce body deformations (except due to rolling and/or sliding friction against the cell cylinder). **F**-**G**. Model for sheath-enforced curvature in the *Leptospira* flagellar filament; most (but not all) of the inherent curvature is due to binding of FcpA and FcpB along the convex side of the core, while FlaA1/2 binding on the concave contributes to residual curvature in the absence of FcpA/FcpB.

FcpA and FcpB are required for maximal stabilization of the wild-type supercoiled conformation of *Leptospira* filaments, but filaments maintain a preference for supercoiling even in the absence of both these components. Despite lacking both FcpA and FcpB, the *fcpA*^−^ filaments are markedly more curved than ‘straight’ filaments from cryo-EM specimens of tobacco mosaic virus filaments or amyloid fibrils (estimated at ~0.7 μm^−1^ when analyzed similarly(Rohou & Grigorieff, 2014)). Unlike the wild-type and *fcpB*^−^ samples, *fcpA*^−^ filament tomogram segments divide into two populations of roughly equal sizes with different diameters (Supplementary Fig. 9); curvature distributions of both populations show signs of heterogeneity, and trajectories from the smaller-diameter population tend towards slightly lesser curvature (Fig. 5C). This latter observation indicates that sheath component(s) other than FcpA and FcpB (perhaps FlaA1and/or FlaA2) may also contribute to filament curvature.

## Discussion

Here we have identified a novel, asymmetric architecture in the *Leptospira* flagellum, previously unobserved either in exoflagella or in other spirochetes. We have further demonstrated that this asymmetry is functionally linked to flagellar supercoiling, a key attribute associated with spirochete motility. An important innovation in the current work was the application of a subtomogram averaging procedure to solve structures of the highly stable, supercoiled form present in wild-type *Leptospira* flagellar filaments. This approach contrasts with previous structural studies of flagellar filaments, almost all of which utilized mutant forms that strongly favor a straight, helically symmetric filament conformation, as was required for high-resolution cryo-EM analysis. To our knowledge, only one prior study reports the structure of a supercoiled spirochete flagellum(Liu et al., 2010); however, in that case the filament was analyzed under the assumption of helical symmetry and was computationally straightened, thus obscuring any asymmetry. Thus, the question of asymmetry in the flagellar filament, in particular for spirochetes, has not been addressed until now.

The asymmetric structure of the *Leptospira* flagellar filament presented here illustrates how curved filament morphology and stiffness– two other critical features for spirochetal motility as indicated by physics modelling studies(Dombrowski et al., 2009; Kan & Wolgemuth, 2007)– can be enforced via the addition of a unique, asymmetric sheath surrounding the filament core. To support the large loads involved in driving whole-body undulations, spirochete endoflagella possess specialized reinforcements, including extra-large motors capable of exerting higher torques than conventional exoflagellar motors(Beeby et al., 2016) and unique hooks exhibiting covalent crosslinking between FlgE subunits(Lynch et al., 2019). Moreover, *Leptospira,* alone among the known spirochetes, has only a single flagellum at each cell end, whereas *Borrelia, Treponema*, and other species exhibit multiple copies per cell end(Charon et al., 2012). One function of the sheath elements FcpA and FcpB, to date found only in *Leptospira*, may therefore be to reinforce the endoflagellum to allow for even higher torque loads required in the absence of additional, supporting flagella.

Our data indicate that the asymmetric outer sheath locks the *Leptospira* flagellar filament into a single, supercoiled conformer, as reflected by a single peak in its curvature distribution (Figure 5). In contrast, filaments from exoflagellated bacteria exhibit structural polymorphism, fluctuating between a multitude of supercoiled states with differing geometries(Maki-Yonekura, Yonekura, & Namba, 2010). Substantially greater flexibility observed in *Leptospira* mutants lacking one or more sheath elements (Fig. 5A-C) may reflect an analogous polymorphic switching function of the filament core, although our data are inconclusive in this regard. In any case, the asymmetric sheath composition identified here in *Leptospira* is ideally suited to trigger flagellar supercoiling, by introducing unequal mechanical distortion on opposite sides of the filament (Fig. 5F-G).

Although FcpA and FcpB play conspicuous roles in supercoiling and asymmetry determination, our images of *fcpA*^−^ filaments indicate that the remaining sheath components are themselves capable of asymmetric binding, and do so preferentially on the concave side of the filament (Fig. 3; Supplementary Fig. 9; Supplementary Table 3). These sheath proteins include FlaA1 and FlaA2, although we are not currently able to localize these in the filament map. Orthologs of *Leptospira* FlaA proteins are found in all spirochetes (Supplementary Table 1). Furthermore, loss of FlaA has been shown to modify the diameter(Li, Corum, et al., 2000) and superhelical pitch(Lambert et al., 2012) of spirochetal filaments. These observations raise the possibility that asymmetric arrangement of components in the sheath (and/or the core) may be a general feature that contributes to supercoiling in spirochete flagella.

## Acknowledgments

We thank the IT Department from Institut Pasteur for protein modeling computations on the TARS cluster and the staff in the Yale High-Performance Computing facility for their maintenance of these facilities. We thank Prof. Felix Rey for providing insightful discussions about the manuscript.

## Author contributions

A.B, A.I.K, J.L., M.P., C.V.S., F.T. and E.A.W.J. conceived the study. M.B., K.H.G, J.L., M., F.S.M., M.P., Z.S., F.T., and E.A.W.J. carried out the experiments and acquired the data. A.B., M.B., K.H.G., and C.V.S. and F.T. designed experiments, analyzed the data, and wrote the manuscript. A.B. and C.V.S. revised the manuscript and coordinated the project. All authors gave final approval for publication.

## Data and Materials Availability

Atomic models for three crystal forms of FcpA, and for FcpB, have been deposited in the Protein Data Bank under accession numbers 6NQW, 6NQX, 6NQY, and 6NQZ (respectively). Cryo-EM maps have been deposited in the Electron Microscopy Data Bank (EMDB) under accession number EMD-20504. A pseudo-atomic model consisting of main chain FlaB, FcpA and FcpB docked into the wild-type filament reconstruction has been deposited in the Protein Data Bank (PDB) under the accession number 6PWB.

## References

Beeby, M., Ribardo, D. A., Brennan, C. A., Ruby, E. G., Jensen, G. J., & Hendrixson, D. R. (2016). Diverse high-torque bacterial flagellar motors assemble wider stator rings using a conserved protein scaffold. Proc Natl Acad Sci U S A, 113(13), E1917–1926. doi:10.1073/pnas.1518952113

Brahamsha, B., & Greenberg, E. P. (1988). Biochemical and cytological analysis of the complex periplasmic flagella from Spirochaeta aurantia. J Bacteriol, 170(9), 4023–4032. doi:10.1128/jb.170.9.4023-4032.1988

Bromley, D. B., & Charon, N. W. (1979). Axial filament involvement in the motility of Leptospira interrogans. J Bacteriol, 137(3), 1406–1412. Retrieved from http://www.ncbi.nlm.nih.gov/pubmed/438121

Carazo, J.-M., Herman, G. T., Sorzano, C. O. S., & Marabini, R. (2006). Algorithm for three-dimensional reconstruction from the imperfect projection data provided by electron microscopy. In J. Frank (Ed.), Electron Tomography: Methods for three-dimensional visualization of structures in the cell (pp. 217–243). New York: Springer.

Charon, N. W., Cockburn, A., Li, C., Liu, J., Miller, K. A., Miller, M. R., …Wolgemuth, C. W. (2012). The unique paradigm of spirochete motility and chemotaxis. Annu Rev Microbiol, 66, 349–370. doi:10.1146/annurev-micro-092611-150145

Charon, N. W., & Goldstein, S. F. (2002). Genetics of motility and chemotaxis of a fascinating group of bacteria: the spirochetes. Annu Rev Genet, 36, 47–73. doi:10.1146/annurev.genet.36.041602.134359

Charon, N. W., Goldstein, S. F., Block, S. M., Curci, K., Ruby, J. D., Kreiling, J. A., & Limberger, R. J. (1992). Morphology and dynamics of protruding spirochete periplasmic flagella. J Bacteriol, 174(3), 832–840. doi:10.1128/jb.174.3.832-840.1992

Dombrowski, C., Kan, W., Motaleb, M. A., Charon, N. W., Goldstein, R. E., & Wolgemuth, C. W. (2009). The elastic basis for the shape of Borrelia burgdorferi. Biophys J, 96(11), 4409–4417. doi:10.1016/j.bpj.2009.02.066

Erhardt, M., Namba, K., & Hughes, K. T. (2010). Bacterial nanomachines: the flagellum and type III injectisome. Cold Spring Harb Perspect Biol, 2(11), a000299. doi:10.1101/cshperspect.a000299

Galkin, V. E., Yu, X., Bielnicki, J., Heuser, J., Ewing, C. P., Guerry, P., & Egelman, E. H. (2008). Divergence of quaternary structures among bacterial flagellar filaments. Science, 320(5874), 382–385. doi:10.1126/science.1155307

Goldstein, S. F., & Charon, N. W. (1988). Motility of the spirochete Leptospira. Cell Motil Cytoskeleton, 9(2), 101–110. doi:10.1002/cm.970090202

Holm, L., & Laakso, L. M. (2016). Dali server update. Nucleic Acids Res, 44(W1), W351–355. doi:10.1093/nar/gkw357

Jiang, J., Zhang, Y., Krainer, A. R., & Xu, R. M. (1999). Crystal structure of human p32, a doughnut-shaped acidic mitochondrial matrix protein. Proc Natl Acad Sci U S A, 96(7), 3572–3577. doi:10.1073/pnas.96.7.3572

Kan, W., & Wolgemuth, C. W. (2007). The shape and dynamics of the Leptospiraceae. Biophys J, 93(1), 54–61. doi:10.1529/biophysj.106.103143

Ko, A. I., Goarant, C., & Picardeau, M. (2009). Leptospira: the dawn of the molecular genetics era for an emerging zoonotic pathogen. Nat Rev Microbiol, 7(10), 736–747. doi:10.1038/nrmicro2208

Lambert, A., Picardeau, M., Haake, D. A., Sermswan, R. W., Srikram, A., Adler, B., & Murray, G. A. (2012). FlaA proteins in Leptospira interrogans are essential for motility and virulence but are not required for formation of the flagellum sheath. Infect Immun, 80(6), 2019–2025. doi:10.1128/IAI.00131-12

Li, C., Corum, L., Morgan, D., Rosey, E. L., Stanton, T. B., & Charon, N. W. (2000). The spirochete FlaA periplasmic flagellar sheath protein impacts flagellar helicity. J Bacteriol, 182(23), 6698–6706. Retrieved from https://www.ncbi.nlm.nih.gov/pubmed/11073915

Li, C., Motaleb, A., Sal, M., Goldstein, S. F., & Charon, N. W. (2000). Spirochete periplasmic flagella and motility. J Mol Microbiol Biotechnol, 2(4), 345–354. Retrieved from https://www.ncbi.nlm.nih.gov/pubmed/11075905

Li, C., Wolgemuth, C. W., Marko, M., Morgan, D. G., & Charon, N. W. (2008). Genetic analysis of spirochete flagellin proteins and their involvement in motility, filament assembly, and flagellar morphology. J Bacteriol, 190(16), 5607–5615. doi:10.1128/JB.00319-08

Liu, J., Howell, J. K., Bradley, S. D., Zheng, Y., Zhou, Z. H., & Norris, S. J. (2010). Cellular architecture of Treponema pallidum: novel flagellum, periplasmic cone, and cell envelope as revealed by cryo electron tomography. J Mol Biol, 403(4), 546–561. doi:10.1016/j.jmb.2010.09.020

Lynch, M. J., Miller, M., James, M., Zhang, S., Zhang, K., Li, C., …Crane, B. R. (2019). Structure and chemistry of lysinoalanine crosslinking in the spirochaete flagella hook. Nat Chem Biol. doi:10.1038/s41589-019-0341-3

Madej, T., Lanczycki, C. J., Zhang, D., Thiessen, P. A., Geer, R. C., Marchler-Bauer, A., & Bryant, S. H. (2014). MMDB and VAST+: tracking structural similarities between macromolecular complexes. Nucleic Acids Res, 42(Database issue), D297–303. doi:10.1093/nar/gkt1208

Maki-Yonekura, S., Yonekura, K., & Namba, K. (2010). Conformational change of flagellin for polymorphic supercoiling of the flagellar filament. Nat Struct Mol Biol, 17(4), 417–422. doi:10.1038/nsmb.1774

McBride, A. J., Athanazio, D. A., Reis, M. G., & Ko, A. I. (2005). Leptospirosis. Curr Opin Infect Dis, 18(5), 376–386. Retrieved from http://www.ncbi.nlm.nih.gov/pubmed/16148523

Motaleb, M. A., Corum, L., Bono, J. L., Elias, A. F., Rosa, P., Samuels, D. S., & Charon, N. W. (2000). Borrelia burgdorferi periplasmic flagella have both skeletal and motility functions. Proc Natl Acad Sci U S A, 97(20), 10899–10904. doi:10.1073/pnas.200221797

Nauman, R. K., Holt, S. C., & Cox, C. D. (1969). Purification, ultrastructure, and composition of axial filaments from Leptospira. J Bacteriol, 98(1), 264–280. Retrieved from https://www.ncbi.nlm.nih.gov/pubmed/4891807

Picardeau, M. (2017). Virulence of the zoonotic agent of leptospirosis: still terra incognita? Nat Rev Microbiol, 15(5), 297–307. doi:10.1038/nrmicro.2017.5

Picardeau, M., Brenot, A., & Saint Girons, I. (2001). First evidence for gene replacement in Leptospira spp. Inactivation of L. biflexa flaB results in non-motile mutants deficient in endoflagella. Mol Microbiol, 40(1), 189–199. Retrieved from https://www.ncbi.nlm.nih.gov/pubmed/11298286

Rohou, A., & Grigorieff, N. (2014). Frealix: model-based refinement of helical filament structures from electron micrographs. J Struct Biol, 186(2), 234–244. doi:10.1016/j.jsb.2014.03.012

San Martin, F., Mechaly, A. E., Larrieux, N., Wunder, E. A., Jr., Ko, A. I., Picardeau, M., …Buschiazzo, A. (2017). Crystallization of FcpA from Leptospira, a novel flagellar protein that is essential for pathogenesis. Acta Crystallogr F Struct Biol Commun, 73(Pt 3), 123–129. doi:10.1107/S2053230X17002096

Sasaki, Y., Kawamoto, A., Tahara, H., Kasuga, K., Sato, R., Ohnishi, M., …Koizumi, N. (2018). Leptospiral flagellar sheath protein FcpA interacts with FlaA2 and FlaB1 in Leptospira biflexa. PLoS One, 13(4), e0194923. doi:10.1371/journal.pone.0194923

Sheikh, M. A., Potter, J. A., Johnson, K. A., Sim, R. B., Boyd, E. F., & Taylor, G. L. (2008). Crystal structure of VC1805, a conserved hypothetical protein from a Vibrio cholerae pathogenicity island, reveals homology to human p32. Proteins, 71(3), 1563–1571. doi:10.1002/prot.21993

Sultan, S. Z., Manne, A., Stewart, P. E., Bestor, A., Rosa, P. A., Charon, N. W., & Motaleb, M. A. (2013). Motility is crucial for the infectious life cycle of Borrelia burgdorferi. Infect Immun, 81(6), 2012–2021. doi:10.1128/IAI.01228-12

Tahara, H., Takabe, K., Sasaki, Y., Kasuga, K., Kawamoto, A., Koizumi, N., & Nakamura, S. (2018). The mechanism of two-phase motility in the spirochete Leptospira: Swimming and crawling. Sci Adv, 4(5), eaar7975. doi:10.1126/sciadv.aar7975

Vernel-Pauillac, F., & Werts, C. (2018). Recent findings related to immune responses against leptospirosis and novel strategies to prevent infection. Microbes Infect, 20(9-10), 578–588. doi:10.1016/j.micinf.2018.02.001

Wang, F., Burrage, A. M., Postel, S., Clark, R. E., Orlova, A., Sundberg, E. J., …Egelman, E. H. (2017). A structural model of flagellar filament switching across multiple bacterial species. Nat Commun, 8(1), 960. doi:10.1038/s41467-017-01075-5

Wolgemuth, C. W. (2015). Flagellar motility of the pathogenic spirochetes. Semin Cell Dev Biol, 46, 104–112. doi:10.1016/j.semcdb.2015.10.015

Wolgemuth, C. W., Charon, N. W., Goldstein, S. F., & Goldstein, R. E. (2006). The flagellar cytoskeleton of the spirochetes. J Mol Microbiol Biotechnol, 11(3-5), 221–227. doi:10.1159/000094056

Wunder, E. A., Jr., Figueira, C. P., Benaroudj, N., Hu, B., Tong, B. A., Trajtenberg, F., …Ko, A. I. (2016). A novel flagellar sheath protein, FcpA, determines filament coiling, translational motility and virulence for the Leptospira spirochete. Mol Microbiol, 101(3), 457–470. doi:10.1111/mmi.13403

Wunder, E. A., Jr., Figueira, C. P., Santos, G. R., Lourdault, K., Matthias, M. A., Vinetz, J. M., …Ko, A. I. (2016). Real-Time PCR Reveals Rapid Dissemination of Leptospira interrogans after Intraperitoneal and Conjunctival Inoculation of Hamsters. Infect Immun, 84(7), 2105–2115. doi:10.1128/IAI.00094-16

Wunder, E. A., Jr., Slamti, L., Suwondo, D. N., Gibson, K. H., Shang, Z., Sindelar, C. V., …Picardeau, M. (2018). FcpB Is a Surface Filament Protein of the Endoflagellum Required for the Motility of the Spirochete Leptospira. Front Cell Infect Microbiol, 8, 130. doi:10.3389/fcimb.2018.00130

Yamashita, I., Hasegawa, K., Suzuki, H., Vonderviszt, F., Mimori-Kiyosue, Y., & Namba, K. (1998). Structure and switching of bacterial flagellar filaments studied by X-ray fiber diffraction. Nat Struct Biol, 5(2), 125–132. Retrieved from https://www.ncbi.nlm.nih.gov/pubmed/9461078

Yonekura, K., Maki-Yonekura, S., & Namba, K. (2003). Complete atomic model of the bacterial flagellar filament by electron cryomicroscopy. Nature, 424(6949), 643–650. doi:10.1038/nature01830

