## Supplementary Data, Methods, Figures, Tables for "An asymmetric sheath controls flagellar supercoiling and motility in the Leptospira spirochete"

**Supplementary Materials:**

### Materials and Methods

**Strains and cell culturing.** *Leptospira biflexa* serovar Patoc strain Patoc I (Paris) wild type, *fcpA*⁻ and *fcpB*- mutant cells were cultured in Ellinghausen–McCullough–Johnson–Harris (EMJH) liquid medium until they reached logarithmic phase at 30°C(Wunder et al., 2016; Wunder et al., 2018).

**Periplasmic flagella purification.** Purification of periplasmic flagella was performed as described.(Wunder et al., 2016; Wunder et al., 2018).

**Recombinant protein crystallization and crystal structure determination.** FcpA (Uniprot B0STJ8) from *L. biflexa* serovar Patoc strain Patoc I (Paris) was expressed, purified and crystallized as previously reported(San Martin et al., 2017). Three different crystal forms were obtained (Supplementary Table 2), and the structure was solved *ab initio* with the hexagonal data set using Arcimboldo(Rodriguez et al., 2009). Resulting electron density maps were automatically traceable(Lamzin, Perrakis, & Wilson, 2012), and the model was subsequently refined with Buster(Bricogne G. & Roversi P, 2017) iterated with manual rebuilding and validation with Coot(Emsley, Lohkamp, Scott, & Cowtan, 2010). The other two monoclinic crystal forms were solved by molecular replacement(McCoy et al., 2007) using the hexagonal structure as a search probe, then further refined following similar procedures. The final refined model spans residues 55 to 291, while the first 54 residues toward the N-terminus were disordered.

The gene LIC11848 (Uniprot Q72RA0) coding for FcpB from *L. interrogans* serovar Copenhageni strain Fiocruz L1-130, was cloned in expression plasmid pQE80 (Qiagen), including a TEV-cleavable 6xHis tag. Overexpression was achieved in *E. coli* Rosetta-gami 2 (DE3), induction conditions, chromatographic purification and TEV cleavage procedures were performed as for FcpA(San Martin et al., 2017). After cleavage, FcpB started at sequence SQQNSGS (thus excluding the signal peptide), with an N-terminal overhanging tripeptide GSG derived from the plasmid. Single FcpB crystals were grown at 20°C using vapor diffusion technique in hanging drops, mixing 2 µl protein solution (18 mg/mL in 20 mM Tris.HCl pH 8.0, 150 mM NaCl) with 2µl reservoir solution 0.4M $NH_4I$, 26% (w/v) PEG 3350, 0.05M MES pH 6.5, 21% glycerol. Initial anomalous signal was detectable only to 4.5Å, improved signal was achieved by quick-soaking crystals in 0.1M "magic triangle" I3C(Beck, Krasauskas, Gruene, & Sheldrick, 2008) for 20 s. Highly redundant data sets were processed using XDS(Kabsch, 2010) and scaled with Aimless(Evans, 2011). The structure was solved by SAD, exploiting the iodine atoms as anomalous scatterers, the iodine sub-structure was solved with ShelxD(Schneider & Sheldrick, 2002) and refined with Sharp(Bricogne, Vonrhein, Flensburg, Schiltz, & Paciorek, 2003). Density modification was performed with Pirate and resulting electron density maps allowed for manual chain tracing and initial model refinement with Buster. A second dataset diffracting X-rays to higher resolution was eventually used for final refinement with Buster, iterated with manual rebuilding and validation with Coot.

Structural analyses were done with the CCP4 suite(Winn et al., 2011) of programs and with Pymol(Schrodinger, 2015).

**Cryo-electron microscopy.** *L. biflexa* wild-type flagella**.** Purified *L. biflexa* wild-type flagella were mixed with 6x concentrated Protein A conjugated with 10 nm colloidal Au (EMS Aurion, Hatfield, PA). Prior to sample application, the 300 mesh Cu Quantifoil grids (Ted Pella, Inc., Redding, CA) with 1.2 µm diameter holes were plasma discharged with $H_2O_2$ for 30 seconds in a Model 950, Solarus Advanced Plasma System (GATAN). Approximately 3 uL of 2:1 (v/v) flagella: 6x Protein A Gold was applied to each grid. The grids were plunge frozen into liquid ethane in a Mark III Vitrobot (FEI Company, Eindhoven, The Netherlands) following a 2-minute incubation time, blot time of 6-7.5 seconds, and blot offset of -2 mm at 18°C and 100% humidity.

Tilt series were acquired on a 300 kV Polara cryo-TEM (UT Houston, Houston, TX) using SerialEM(Mastronarde, 2005) (University of Colorado, Boulder, CO) equipped with a K2 direct electron detector (GATAN, Inc., Pleasanton, CA). Tilt series were acquired with a defocus of between -2.0 and -4.0 µm at +/- 53° with 3° increments in counting mode at a dose rate of ~8 $e^-$/pixel/second. At each tilt angle, 12 frames were collected with 0.1 s exposure per frame. The total dosage for each tilt series was ~60 $e^-/Å^2$. The pixel size at a magnification of 15500x was calculated to be 2.604 Å at a binning of 1. A total of 120 tilt series were acquired.

*L. biflexa* $fcpA^-$ and $fcpB^-$ flagella**.** Purified $fcpA^-$ and, $fcpB^-$ flagella were prepared similarly to the wild-type flagella and vitrified as described above. Mutant $fcpA^-$ or, separately, $fcpB^-$ flagella were mixed with 6x concentrated 10 nm colloidal Au BSA Gold Tracer beads (EMS Aurion) prior to plunge freezing. The 200 mesh Cu C-flat (EMS) grids with 1.2 µm hole spacing had a 3 nm thick layer of carbon applied to ensure distribution of gold beads and flagella. The grids were plasma discharged with $ArO_2$ for six seconds in a Solaris Advanced Plasma System, Model 950 (GATAN).Tilt series were then acquired on a 200 kV Tecnai F20 cryo-TEM (CCMI, Yale University, New Haven, CT) using SerialEM(Mastronarde, 2005) (Boulder, CO) and imaged with a K2 direct electron detector (GATAN) in counting mode. The Tilt series were acquired over +/- 60° in 3° increments with a defocus of -2.5 µm. At a magnification of 14500x at spot size 7, the dose rate was ~8 $e^-$/pixel/second. The exposure time per tilt angle was 1.5 s with ~2.25 $e^-/Å^2$ and was dose fractionated into 7 frames, each receiving 0.2 s of exposure time. The final dose was ~90 $e^-/Å^2$ with a pixel size was 2.49 Å. A total of 26 tilt series were acquired.

**Tilt series reconstruction and subtomogram averaging.** *L. biflexa* wild-type flagella**.** The frames in each tilt angle in a tilt series were motion corrected using IMOD alignframes(Mastronarde & Held, 2017). The tilt series were then aligned in IMOD, version 4.9.4_RHEL6-64_CUDA8.0(Mastronarde & Held, 2017) using local alignment with the fiduciary markers. After alignment, tilt series underwent CTF-estimation and correction using phase flipping in IMOD, followed by gold subtraction(Himes & Zhang, 2018; Mastronarde & Held, 2017). Final CTF-corrected and aligned tilt series binned by 2, resulting in a pixel size of 5.208 Å and then reconstructed in Tomo3D(Agulleiro & Fernandez, 2011) using weighted back projection. IMOD 3dmod was used to trace a path of particle points along the center of each filament. The addModPts program in PEET(Cope, Heumann, & Hoenger, 2011) was used to set the repeat spacing between each particle point to 52 Å. Using the IMOD model2point program, the particle model file was converted to a text file format that designated X, Y, and Z coordinates for each filament and assigned an incrementing number for each individual filament for import into RELION version 2.1.b1-gcccuda-2016.10-cc37(Scheres, 2012).

For sub-tomogram averaging, the RELION software package(Scheres, 2012) was used for initial sub-tomogram alignment, emClarity(Himes & Zhang, 2018) for high-resolution refinement and 3D reconstruction, and an in-house method for smoothing and interpolation of sub-volume coordinates(Huehn et al., 2018) (Supplementary Fig. 4).

A variation of the RELION pre-processing PYTHON script(Bharat, Russo, Lowe, Passmore, & Scheres, 2015) was developed in house to sort particles by filament to ensure splitting of particles for gold standard FSC calculation. This ensured that two adjacent particles in a filament would not be randomly sorted into separate groups, thereby avoiding potential particle coordinate overlap if two or more adjacent particles occupied the same repeat within a filament during averaging and alignment. To ensure minimal polarity switching during particle alignment, larger cubed segments of filaments (500 Å) were extracted and CTF-corrected using CTFFind4. The resulting alignment parameters were imported into RELION version 2.1 beta for unsupervised sub-tomogram averaging using RELION 3D auto-refinement. No mask was used. For 3D auto-refinement, particle dimensions were set to 490 Å to include 490/52=~9.4 subunits along the filament, and the particle symmetry was designated as C1 due to the inherent asymmetry of the filament. Helical reconstruction options are not implemented for subtomogram averaging in this version of RELION, so the refinement was performed in single-particle mode. An initial angular search spacing of 15° was used. A local search threshold of 1.8° were used for auto-sampling. For local searches, the angular search range in psi and theta (tilt) were 10° and 15°, respectively. The final averaged volume was reported to be 18 Å after FSC calculation during RELION post-processing. Subtomogram averaging in RELION utilized a total of the best 62 tomograms with 10,851 particles.

The flagellum filament structure resulting from subvolume averaging in RELION had a final resolution of 18 Å, but evidently suffered from severe reconstruction artifacts. Efforts to use the RELION subvolume average as a search template for the emClarity(Himes & Zhang, 2018) particle picking procedure failed to yield useful particle coordinates. In-house scripts were therefore utilized to export the RELION alignment parameters to emClarity. Following coordinate import, CTF estimation and correction was performed in emClarity. The optimal defocus estimate was set to -3.0 µm and the defocus window was 0.5 µm.

For averaging and alignment steps in emClarity, particle dimensions and box sizes, as well as in-built shape masks were specified in the emClarity parameter file. Following initial estimates for a filament diameter of ~240 Å and a repeat spacing of 52 Å, the particle 'radius' was set to (145 Å, 145 Å, 26 Å) in the parameter file (param0.m). A rectangular alignment mask with x, y, z dimensions of 200, 200, and 160 Å was applied, encompassing 6-8 repeats. Anisotropic SSNR calculation was activated (flgCones=1). To avoid FSC overestimation due to mixing of closely packed particles together between half datasets, the fscGoldSplitOnTomos option was also activated.

Iterative averaging and alignment steps were run in emClarity for 3-5 cycles with increasingly restrictive search angles and translational shifts. Particle coordinates and Euler angles from several cycles (1, 3 and 5) were extracted from the emClarity results file for analysis of individual filaments. Areas in the tomograms where filaments had lower SNR with respect to the background solvent had poorer tracking of particle coordinates along the filament trajectory, leading to 'breaks' and translational shifts in the x,y,z coordinates or Euler angle misalignments. For each filament, particle coordinates were extracted and graphed using in-house scripts. Particle outliers were deleted, and an in-house script was used to smooth a trajectory through the remaining coordinates, and to fill in gaps in the smoothed trajectory with particles according to a repeat spacing of 52 Å. The particle assessment, excision, and trajectory smoothing procedures were performed several times until the particle coordinates followed a regular, smoothed trajectory. These final coordinates were then re-imported into emClarity for a final 8 cycles of averaging and alignment at a binning of 2 for cycles 0 through 3, and a binning of 1 for cycles 4 through 8. The angular search range (rawAngleSearch=1$^{st}$,2$^{nd}$,3$^{rd}$,4$^{th}$,5$^{th}$) can be set for each alignment cycle with the first two values corresponding to the angular range and increment of the out-of-plane search. The 3$^{rd}$ and 4$^{th}$ values correspond to the angular range and increment for the in-plane search. The 5$^{th}$ value was set to 0 to turn off the helical search, which was not effective for our system. At a binning of 2, the rawAngleSearch range for cycles zero through three, in order, were (0,0,7,1,0), (4,1,0,0,0), (0,0,3,0.75,0), and (2,1,0,0,0).For cycles four through eight, at a binning of 1, the rawAngleSearch ranges were (0,0,5,1,0), (3,1,0,0,0), (0,0,3,0.75,0), (2.25,0.75,0,0,0), and (0,0,1.5,0.5,0). The two half-dataset volumes were combined with a B-factor of 0, and the Gold standard FSC at 0.143 was calculated to be 9.83 A with anisotropic resolutions ranging from 8.89-16.17.

*L. biflexa fcpA$^-$* and *fcpB$^-$* flagella**.** A similar sub-tomogram averaging procedure as described above was followed for the *fcpA$^-$* and *fcpB$^-$* flagella tomograms. Specifically, IMOD was used for motion correction (using the program alignframes) and the tilt series were locally aligned through use of the fiduciary markers. CTF correction was also performed in IMOD through phase flipping, followed by subtraction of the fiduciary markers from the tomograms. The final aligned tomograms were binned by 2, resulting in a pixel size of 4.97Å. Tomograms were then reconstructed in Tomo3D using weighted back projection, and 3dmod (IMOD) was used to select the filaments. The addModPts program in PEET was used to add points corresponding to the repeat spacing of 52Å, and the IMOD program model2point was used to prepare the files for import into RELION.

The *fcpB$^-$* tomograms were imported into RELION using the same scripts as described above, with a box size of 480Å. A total of 24 tomograms were imported, corresponding to 5917 particles. In the program 3dautorefine, no initial reference model was used, and the symmetry was C1, the particle mask was 470Å, and the same angular search parameters as for the wild-type structure were used. The resultant *fcpB$^-$* structure had a resolution of 26.5Å, and similar to the wild-type reconstruction, appeared to have large missing wedge artifacts. For *fcpA$^-$*, a total of 25 tomograms were used, yielding 4327 particles; particles were imported into RELION and refined using similar parameters. Due to limited signal quality and/or sample heterogeneity, the *fcpA$^-$* refinement did not yield a converged structure, with inconsistent particle axial rotations along single filaments. Nevertheless, the resulting *fcpA$^-$* subunit x, y, z coordinates yielded continuous trajectories for many of the filaments. Using 2D classification, two distinct *fcpA$^-$* particle diameters were identified, with 2958 particles in a smaller-diameter class (~120Å) and 323 particles in a larger-diameter class (~170Å). Particle coordinates from these two classes were used for subsequent trajectory/curvature analysis.

For the *fcpB*⁻ dataset, the RELION-refined x, y, z coordinates and Euler angles of all 24 tomograms and 5917 particles were imported into emClarity, using the same in-house scripts as described above. CTF correction and estimation was performed, with an initial defocus estimate and window of -5.0 µm and 2.0 µm respectively. Due to reduced signal to noise ratio in the *fcpB*⁻ data, the reference alignment procedure failed to unambiguously establish the polarity of many of the filaments, which were therefore discarded. After this analysis, 1163 particles corresponding to 14 filaments from 10 tomograms remained.

Initial emClarity parameters designated the particle radius in x, y, and z as 150, 150, and 20Å, and applied a rectangular alignment mask of 180,180, 250 Å. As in the wild-type dataset, the flgCones and fscGoldSplitonTomos options were activated. Five cycles of averaging and alignment were used, in a manner similar to that described above, with each cycle using a binning of 2. The rawAngleSearch range for the cycles, in order, were (0,0,7,1,0), (5,1,0,0,0), (0,0,5,1,0), (4,1,0,0,0), (0,0,3,0.75,0), and (0.75,3,0,0,0). After the fifth cycle, the half data sets were combined, giving an average FSC of 18.41 Å, with a range from 14.58Å to 31.67Å at a B-factor of 0.

**Model building, fitting of protein components into tomographic maps and minimization of the *L. biflexa* flagellar filament.** Models of *L. biflexa* FlaB1 and FcpB were built by homology modeling with Rosetta[63]. 10000 models of FlaB1 were thus generated using the structure of the flagellar filament of *B. subtilis* (PDB 5WJT) in a straight symmetric context. In the case of FcpB, we used our *L. interrogans* experimental structure (PDB 6NQZ) as a template, to generate 20000 models.

Using SITUS (Kovacs, Galkin, & Wriggers, 2018; Wriggers, 2012), the averaged volume of the *L. biflexa* wild-type flagella generated by emClarity was converted to a SITUS readable file using the SITUS map2map program. Using the colores program, single monomers of FcpA and, separately, FcpB were rigidly fit into the ~10 Å resolution 3D volume with Laplacian filtering using an angular step of 5° where quasi-uniform angular spacing was enforced by the pole sparsing method. The highest-scoring fittings thus obtained gave alignments for two protofilaments of FcpA and one protofilament of FcpB within the outer convex sheath region. To identify additional docking locations within the sheath where resolution anisotropy (due to the missing cone of Fourier data) may have reduced the efficacy of the fitting procedure, we performed additional SITUS searches with the same parameters as before but with modified input maps and models. Specifically, we (i) focused searches on specific sheath regions by segmenting the cryo-EM map into a series of overlapping sub-regions and (ii) increased the signal power of the FcpA and FcpB search models by generating symmetry-related 'triplexes' of the initially discovered docking hits. Sub-region segmentation of the flagellar sheath in our 3D volume was performed using the UCSF-Chimera(Pettersen et al., 2004) module Segger. To generate a 'triplex' of FcpA or FcpB atomic coordinates, the highest-scoring initial hit was axially shifted by one 52 Å helical repeat forward and backwards to generate a composite PDB file containing three FcpA (or FcpB) monomers at consecutive sites along a single protofilament. Top-scoring hits from this second round of SITUS searching identified four additional protofilaments of FcpA, for a total of 6 protofilaments, and three additional protofilaments of FcpB for a total four protofilaments.

The relative positioning of FcpA and FcpB in our sheath model was further supported by an independent computational docking strategy, in which we utilized pre-formed FcpA/FcpB dimers predicted by computational methods. In order to construct FcpA:FcpB complex, we first modelled *L. biflexa* FcpB using our *L. interrogans* experimental structure (PDB 6NQZ) as a template, using standard homology modelling procedures from Rosetta(Das & Baker, 2008). Through this procedure, more than 20000 models of FcpB were generated, including loops β3β4 and β7β8, absent in the experimental structure because of poor electron density in those regions. The best 100 FcpB models, according to the Rosetta score, were used as input for an exhaustive full docking procedure with our FcpA X-ray model. The best 5000 docked configurations, according to the binding energy score, were rigidly fit into a segmented portion of the electron density map of the flagellar filament using SITUS colores and ranked according to the correlation coefficient. The highest-scoring solutions from this search clustered in two distinct molecular arrangements, one of which yielded a fair approximation of a single FcpA/FcpB heterodimer from our full FcpA/FcpB sheath model.

**Curved FlaB core filament building and fitting.** To build a complete atomic model of the curved filament, an 11-mer of modeled FlaB subunits (co-assembled according to (straight) helical symmetry) was first fit into the pseudo-helically symmetrized core volume. The 11-mer was then deformed to follow a curved path that matched the measured curvature of the tomographically averaged wild-type filament, using UCSF Chimera(Pettersen et al., 2004). This core model, corresponding to a single 52Å repeat of the filament, was then merged with the best SITUS solutions for FcpA (6 protofilaments) and FcpB (4 protofilaments). The resulting core-sheath atomic model, representing a single 52Å filament repeat, was then replicated 7 times along the same curved path to generate an assembly representing 8 52Å repeats. To refine this filament model and eliminate minor clashes, all the protein monomers (protomers) placed in density were subjected to iterative cycles of rigid body, side-chain and backbone minimization using positional and conformational constraints, restrained within the cryoET volume map using a customized protocol in Rosetta(Das & Baker, 2008).

**Curvature Analysis.** Curvature was estimated for each reconstructed filament segment by comparing the 3D coordinates of neighboring filament subunits, and the resulting curvature values summed in histograms. The curvature of the purified flagella was determined in the manner of Crenshaw et al.(Crenshaw, Ciampaglio, & McHenry, 2000), utilizing the smoothed three-dimensional coordinates of the filaments. Consider consecutive three-dimensional points $P_1, P_2, P_3, \ldots, P_N$, with each point spaced 52Å apart (corresponding to the repeat spacing of the filament). Let $C=P_i-P_{(i-10)}$ and $D=P_{(i+10)}-P_i$ from i=11 to i=(N-10); a spacing of 10 is used to minimize noise that might result from minor deviations between consecutive points. Then, curvature (κ) at a particular point can be determined by:

$$\kappa_i^* = \left(\frac{C \cdot D}{|C||D|}\right)\left(\frac{2}{|C| + |D|}\right)$$

The final curvature value is then found by averaging the curvature at successive points, in the following manner:

$$\kappa_{<i>}^* = \frac{\kappa_i^* + \kappa_{i+1}^*}{2}$$

The resultant values are multiplied by 1/(pixel size * binning), giving a curvature with units of 1/Å for points 11 to N-10 along the filament. These values were calculated for the 84 filaments of wild-type that were used in the emClarity analysis, the 14 filaments of the *fcpB*$^-$ sample that were used in the emClarity analysis, and 8 filaments of the *fcpA*$^-$ sample. Due to the small size of the dataset and sample heterogeneity, a high-resolution structure of the *fcpA*$^-$ sample could not be determined.

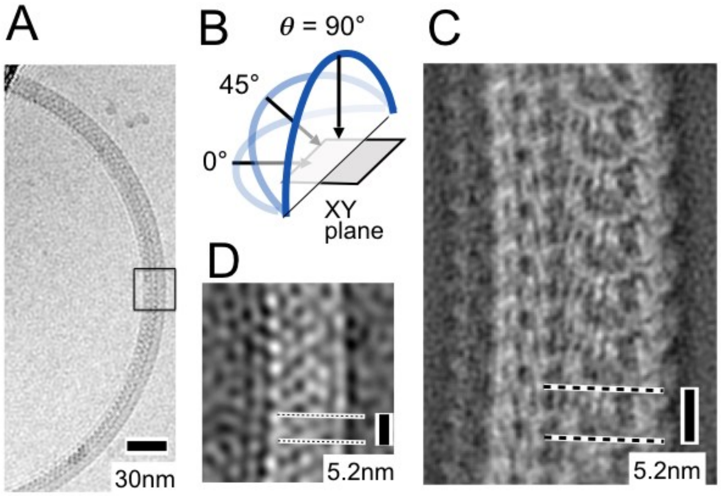


**Supplementary Figure 1** (related to Figure 1). Preferred orientation of purified coiled *L. biflexa* flagellar filaments in the specimen ice layer. **A**. Cryo-electron micrograph showing a wild-type *L. biflexa* flagellar filament. **B**. Schematic illustrating our definition of the tilt angle (***θ***) of a filament with respect to the XY-plane. **C**. Representative 2D class average image from ~250 filaments imaged by cryo-EM at 0° tilt. Within cryo-EM samples, Leptospira flagella are largely restricted to the XY plane (***θ*** ~ 0°) due to confinement within the specimen ice layer, yielding a relatively small number of distinct class average images (< 30). This restriction in the range of ***θ*** values prevents a meaningful 3D reconstruction from being obtained. **D**. Close-up of the boxed region in A after applying a low-pass filter.

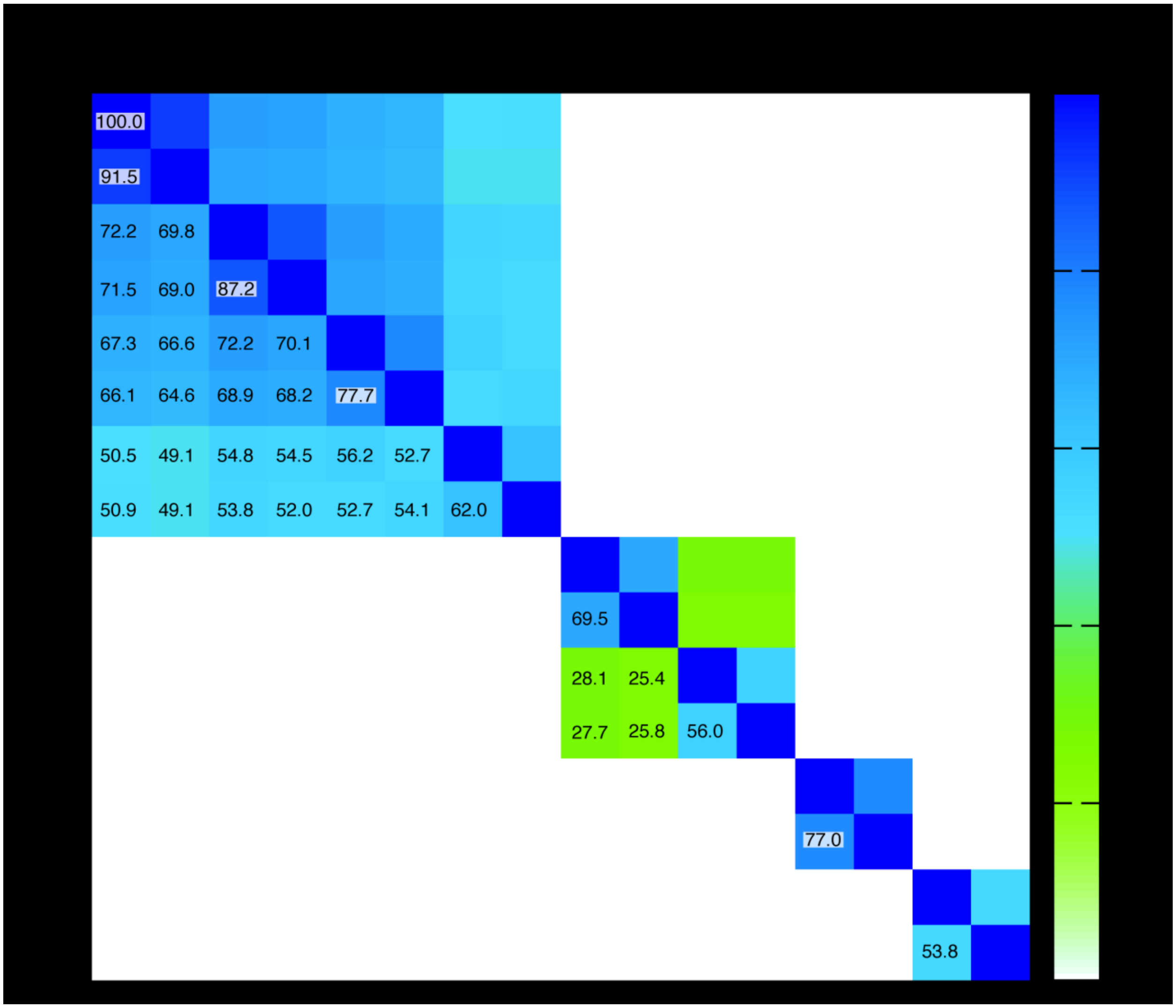


**Supplementary Figure 2** (related to Figure 1). Square matrix plot illustrating pairwise sequence identities of flagellar filament proteins from *Leptospira*. Individual boxes are colored according to the ramp scale shown on the right (white - green - blue). Percent identities are also indicated in each pairwise box (the main diagonal boxes correspond to 100 % self-identities). The prefixes "i" or "b" on the column and row labels indicate *Leptospira interrogans* or *L. biflexa*, respectively, followed by the name of the protein. Note that FlaB3 is significantly less homologous to the other three FlaB isoforms. The two FlaA isoforms, FlaA1 and FlaA2 are homologous, albeit with low sequence identity (<30%). FcpA and FcpB are not homologous between them, and FcpA shows significantly higher sequence conservation among different *Leptospira* species, than FcpB.

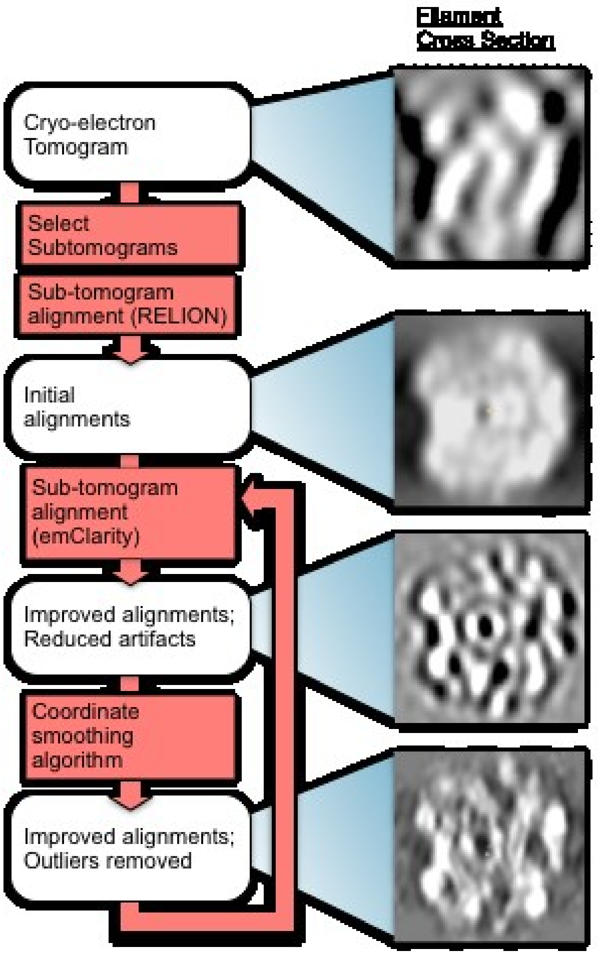


**Supplementary Figure 3** (related to Figure 2). Methods flow chart. Cross-sections of subtomograms or subtomogram averages resulting from each step are shown on the right.

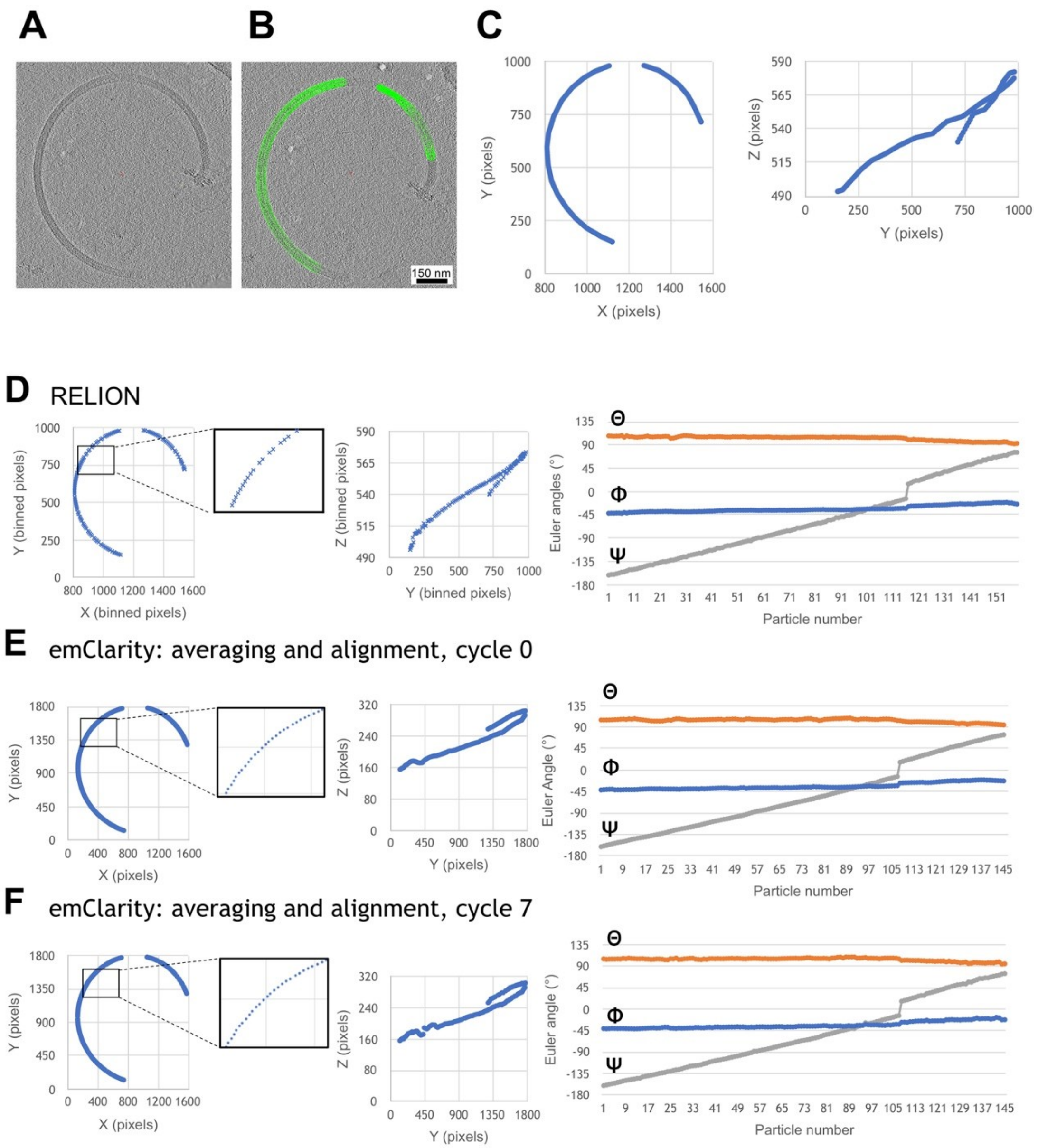


**Supplementary Figure 4** (related to Figure 2). Progressive improvement of alignment parameters for a representative filament, after various stages of refinement. The refinement procedure incorporated an in-house smoothing algorithm that rectifies gaps, duplicates and outliers to obtain continuous 3D coordinate models of every subunit from each selected filament segment. **A**, Section through a reconstructed tomogram of a purified *L. biflexa* WT flagellar filament. **B**, Particle selection (green spheres) that selects the filament trajectory through the tomogram slice. **C**, Particle X/Y (left) and Y/Z (right) coordinates selected in B. Tomograms and coordinates shown here correspond to a binning of 2, at 5.208 Å/pixel. **D.** Filament trajectory (XYZ coordinates and Euler angles) following the initial RELION refinement step. **E**. Filament trajectory output following the first cycle of emClarity refinement. Application of the smoothing algorithm after the RELION step eliminates gaps seen in some sections of the filament trace (compare insets in panels **D** and **E**). **F**. Filament trajectory following an additional smoothing step, followed by a second pair of emClarity/smoothing steps.

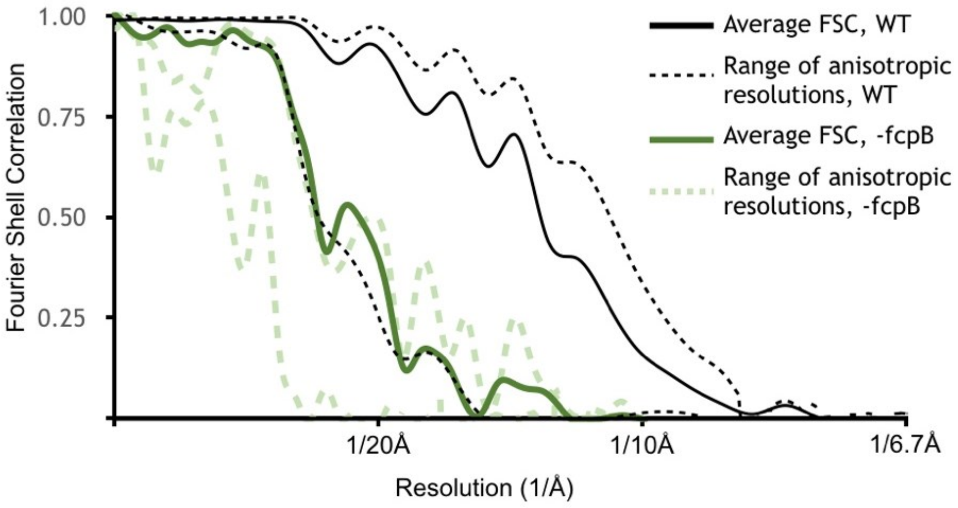


**Supplementary Figure 5** (related to Figure 2). Resolution estimates for wild-type and $fcpB^{-}$ subtomogram average reconstructions. Shown in dashed lines are Fourier shell correlation (FSC) curves that capture the resolution anisotropy using local sectors ('cones') in Fourier space. The corresponding resolution estimates ranged from ~9Å in the best directions (perpendicular to the filament supercoiling axis, i.e. directions parallel to specimen ice layer plane) to ~18Å in the worst direction (parallel to filament supercoiling axis, i.e. a vector perpendicular to the specimen ice layer plane). The latter direction corresponds to the 'missing cone' in our data due to a combination of strongly preferred filament orientation and limited tilt angle in the tomographic data collections. Resolution anisotropy resulted in a marked elongation of map features in this direction (orthogonal to the viewing plane in Fig. 2A, B). Only the highest and lowest-resolution FSC cones are shown.

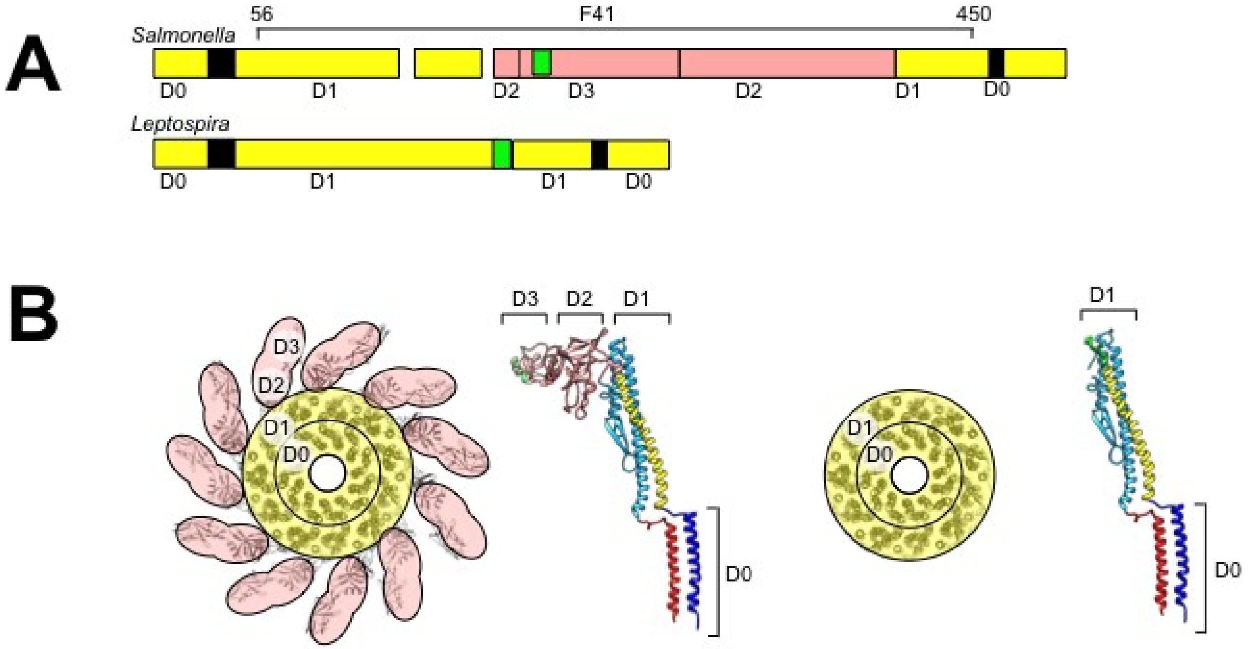


**Supplementary Figure 6** (related to Figure 3). Structure model of *Leptospira* FlaB by homology modeling and sequence alignment. **A.** Schematic of aligned sequences of *Salmonella* FliC (top) and *Leptospira* FlaB (bottom), indicating assignments for subdomains D0 and D1. **B.** Axial view of the *Salmonella* flagellar filament structure (left) and side view of the component FliC structure (right; PDB ID 1UCU[(Yonekura, Maki-Yonekura, & Namba, 2003)]) depicting the locations of subdomains D0-D3. **C.** Corresponding views of the modeled *Leptospira* FlaB core assembly.

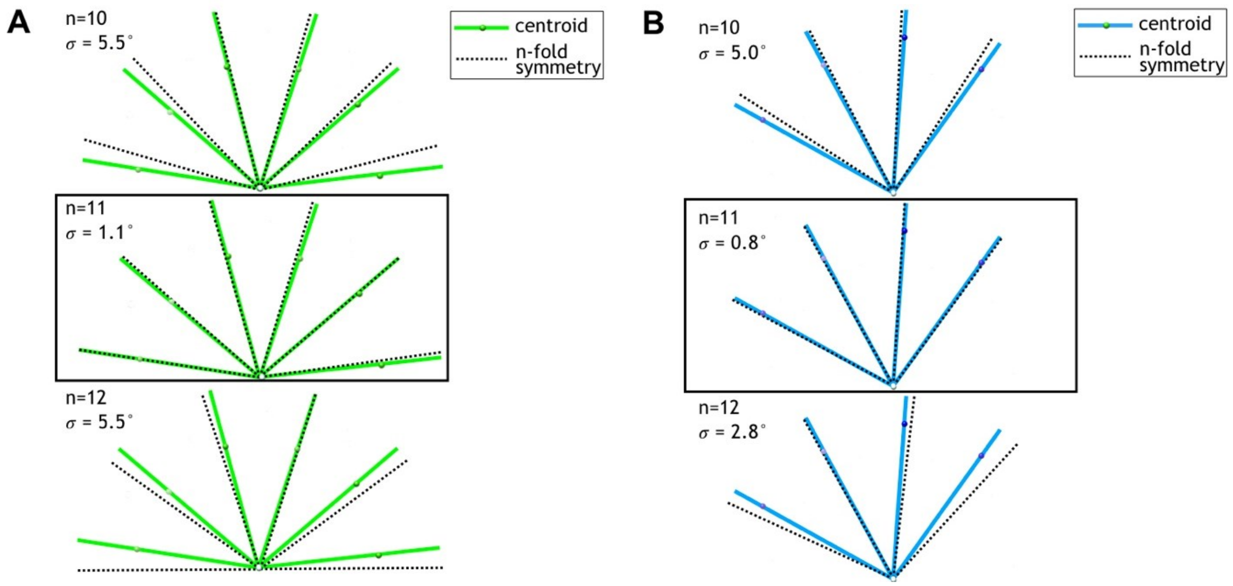


**Supplementary Figure 7** (related to Figure 3). Centroid angular positions of fitted FcpA and FcpB models match an 11-protofilament pseudo-helical lattice. **A.** Filament cross-sectional view showing centroid positions of fitted FcpA models (green spheres) and their angular position (green lines) with respect to the middle of the central channel (white sphere). Overlaid are predicted angular positions (black dashed lines) for symmetric helical lattice sites of the given symmetry type (n=10, 11, or 12 protofilaments). Root mean squared angular deviations between centroid and symmetric angular positions (σ) are smallest for the 11-protofilament case. **B.** Plots of FcpB centroid positions, as in A, indicating best agreement with an 11-protofilament binding pattern.

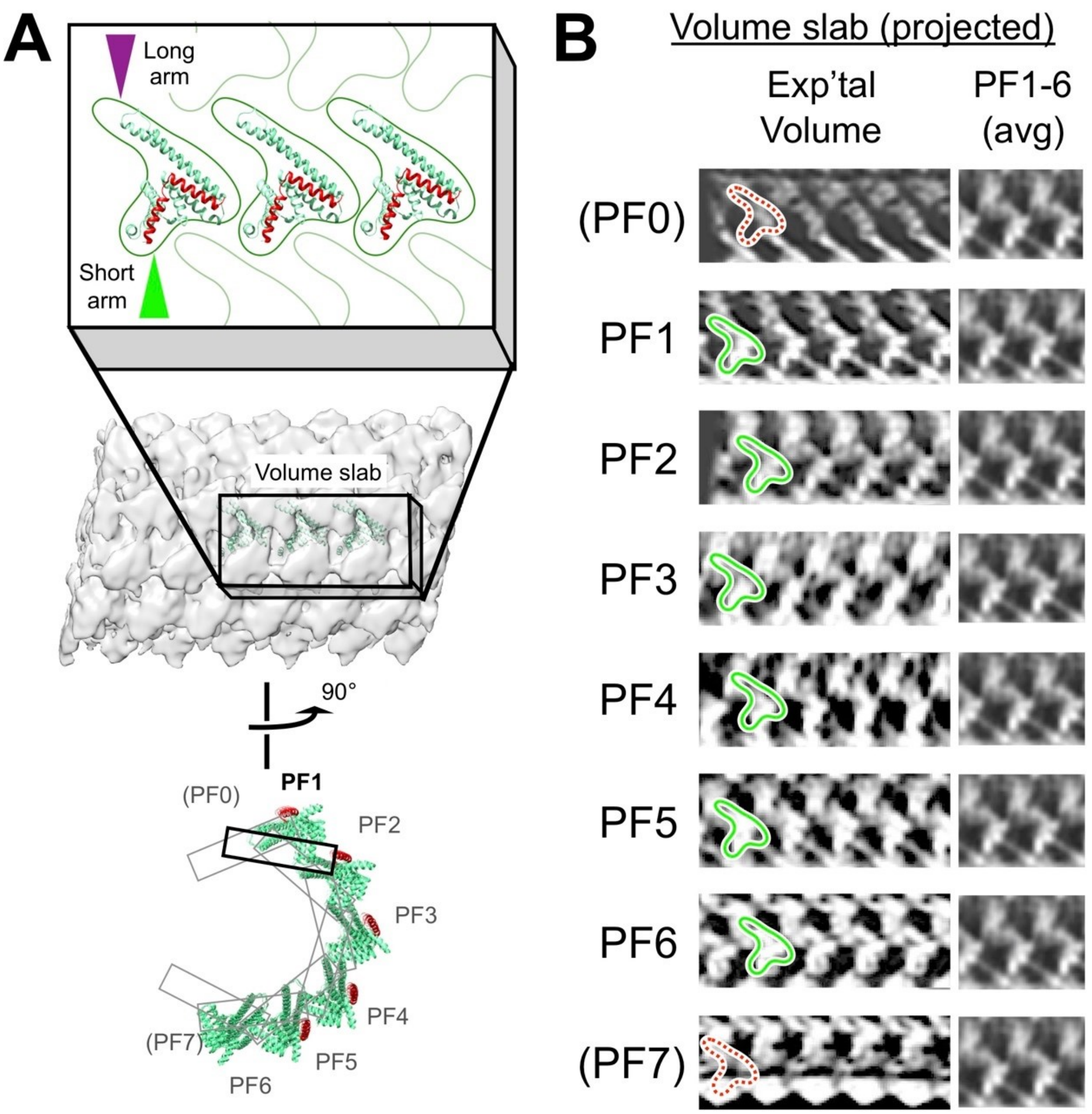


**Supplementary Figure 8** (related to Figure 3). Identification of six similarly arranged FcpA protofilaments in the flagellar filament sheath. **A.** Schematic depicting locations of symmetry-related 'slab' volumes defined within the filament sheath layer. Volume slabs are rotated in progressive increments of 360°/11 = 32.7° about an axis tangent to a curve running through the filament center (which corresponds to the filament helical axis, for a straightened filament). **B.** Two-dimensional projections of the slab volumes in A, revealing six rows ('PF1' – 'PF6') of distinctive 'V'-shaped features consistent with the size and shape of FcpA monomers (green outlines) docked on the core surface. V-shaped features are absent from symmetry-related locations in the first and last volume slabs ('PF0' and 'PF7'), indicating discontinuities in the corresponding pseudo-helical array of FcpA molecules identified by computational docking studies (Fig. 3).

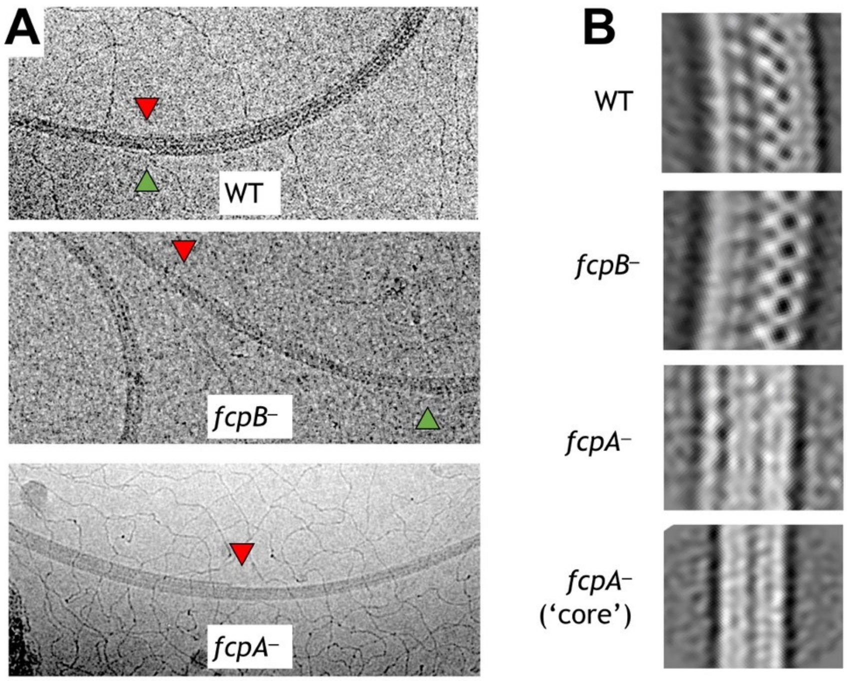


**Supplementary Figure 9** (related to Figure 5). **A.** Cryo-EM images of wild-type and mutant *L. biflexa* flagellar filaments that exhibit shedding of sheath layers, either on the inner (red triangles) or outer (green triangles) curvature. Shedding was only rarely seen in wild-type and *fcpB*$^{-}$ filaments (Supplementary Table 3) and could be observed on both inner and outer curvatures, sometimes in the same filament. In contrast, *fcpA*$^{-}$ filaments usually lost sheath layers from the inner curvature and concurrent shedding on both inner and outer curvatures in this mutant was extremely rare (Supplementary Table 3). **B.** 2D class averaging of filament subtomogram segments reveals distinct filament diameters and/or curvature for wild-type *vs*. mutant flagella. Wild-type flagella subtomogram 2D classes mainly yielded images with a diameter of ~250Å. Subtomogram 2D classes from *fcpB*$^{-}$ samples have a smaller diameter (~95% of wild-type for the dominant class), while classes from *fcpA*$^{-}$ samples are narrower still (66% – 87% of wild-type). Multiple classes of different radii were observed in all samples wild-type and mutant; only the most common ones are shown. Curvature is most evident in the wild-type and *fcpB*$^{-}$ samples, while *fcpA*$^{-}$ filament class averages are straighter; the narrowest class (bottom panel), which likely represents the core FlaB assembly absent a sheath, shows little or no evidence of curvature. Classes similar to the bottom panel were also observed in the wild-type and *fcpB*$^{-}$ samples.

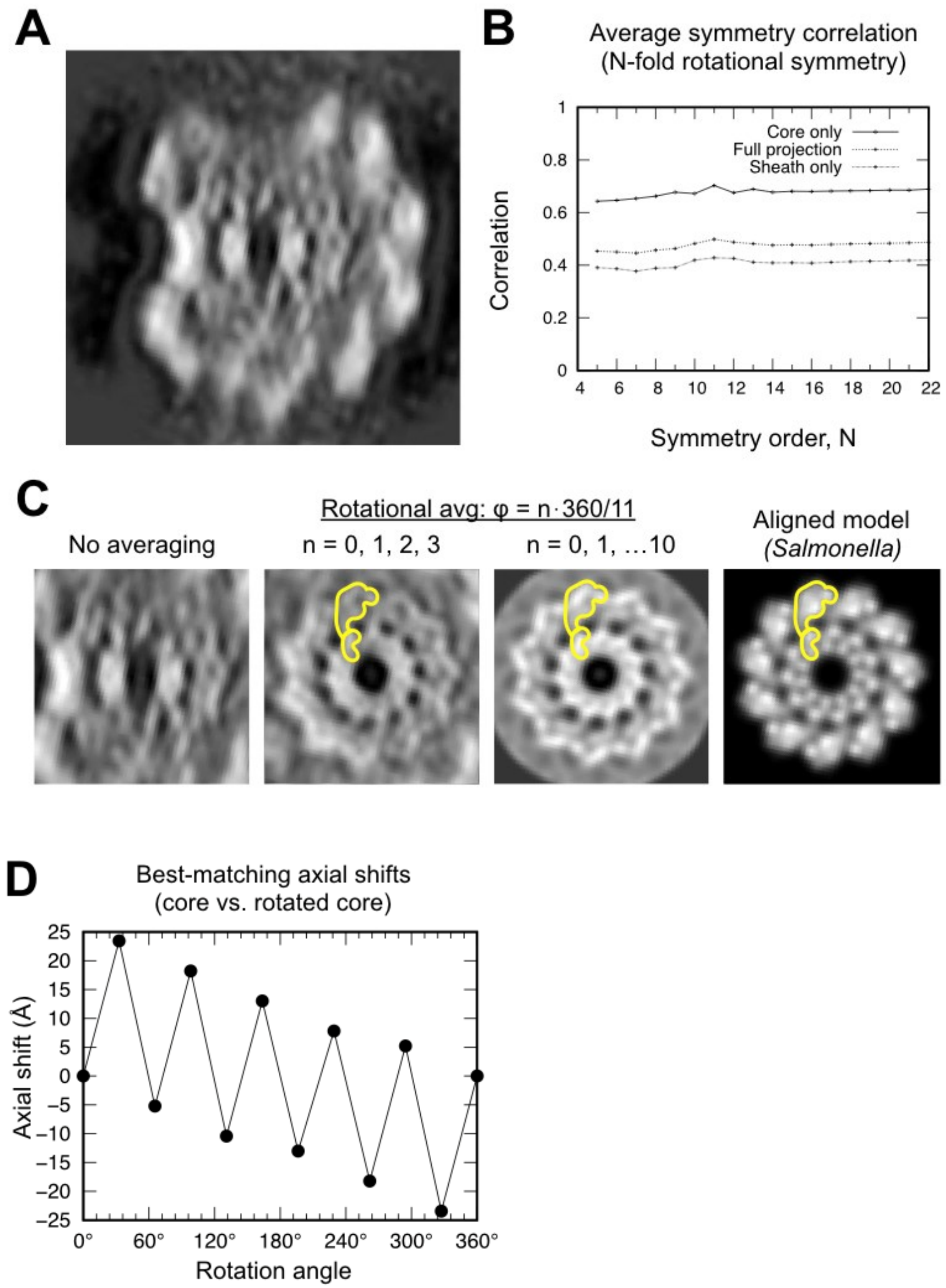


**Supplementary Figure 10** (related to Figure 3). Symmetry analysis of the *Leptospira* core indicates an 11-protofilament architecture. **A.** Projected filament cross section of a 52Å-long segment of the wild-type *L. Biflexa* flagellar filament. **B.** Plot of the averaged cross-correlation between the image in A and *N* rotated copies of itself, corresponding to rotations of 1*360°/*N*, 2*360°/*N*, … (*N*-1)*360°/*N* about the filament center. Thus, for an image containing an 11-fold symmetric feature, the averaged cross-correlation value will be highest for symmetry order *N*=11. Systematically varying N in this calculation reveals a maximum corresponding to 11-fold radial symmetry, matching the symmetry of the *Salmonella* flagellum. Cross-correlations were computed for the entire image ('full projection') as well as masked sub-regions corresponding to the core ('core only') and sheath ('sheath only'). All three of these calculations yield a maximum score for *N*=11. **C.** The 11-fold symmetry operator identified in **B** was used to average the projected map, reducing the effects of the missing wedge and substantially improving molecular features. Leftmost panel shows the original core region, center-left panel shows the result of adding four symmetry-related copies (N=11; $\varphi$ = 0*360°/11, 1*360°/11, 2*360°/11, 3*360°/11) and center-right panel shows the result of adding 11 symmetry-related copies (resulting in an 11-fold symmetric image). Features in the averaged images (yellow shape) resemble the projected D0/D1 subdomain within a projected *Salmonella* flagellar filament cross section (rightmost panel; synthetic image derived from PDB ID 3A5X). **D.** Results of three-dimensional cross-correlation analysis between the wild-type sub-tomographic average volume and rotated copies of itself. For each rotation value ($\varphi$ = 0*360°/11, 1*360°/11, … 11*360°/11), a volume copy was rotated about an axis running through the center channel and masked to exclude all but the core region of a single 52Å axial repeat. A 3D cross-correlation map was then computed between the resulting volume and the original reference, and the axial shift described by the top-scoring peak was plotted for each $\varphi$ rotation value. The resulting graph describes a staggered pattern of helical subunit positions closely matching the 11-start helical symmetry observed in several other reported bacterial flagella structures(Namba, Yamashita, & Vonderviszt, 1989; Wang et al., 2017). The estimated pseudo-helical parameters (~26Å helical pitch, ~5.5 subunits per turn) closely match helical parameters established for several other flagellar filaments (i.e. *Salmonella*: ~25.5Å – 27Å helical pitch, 5.4x – 5.5x subunits per turn).

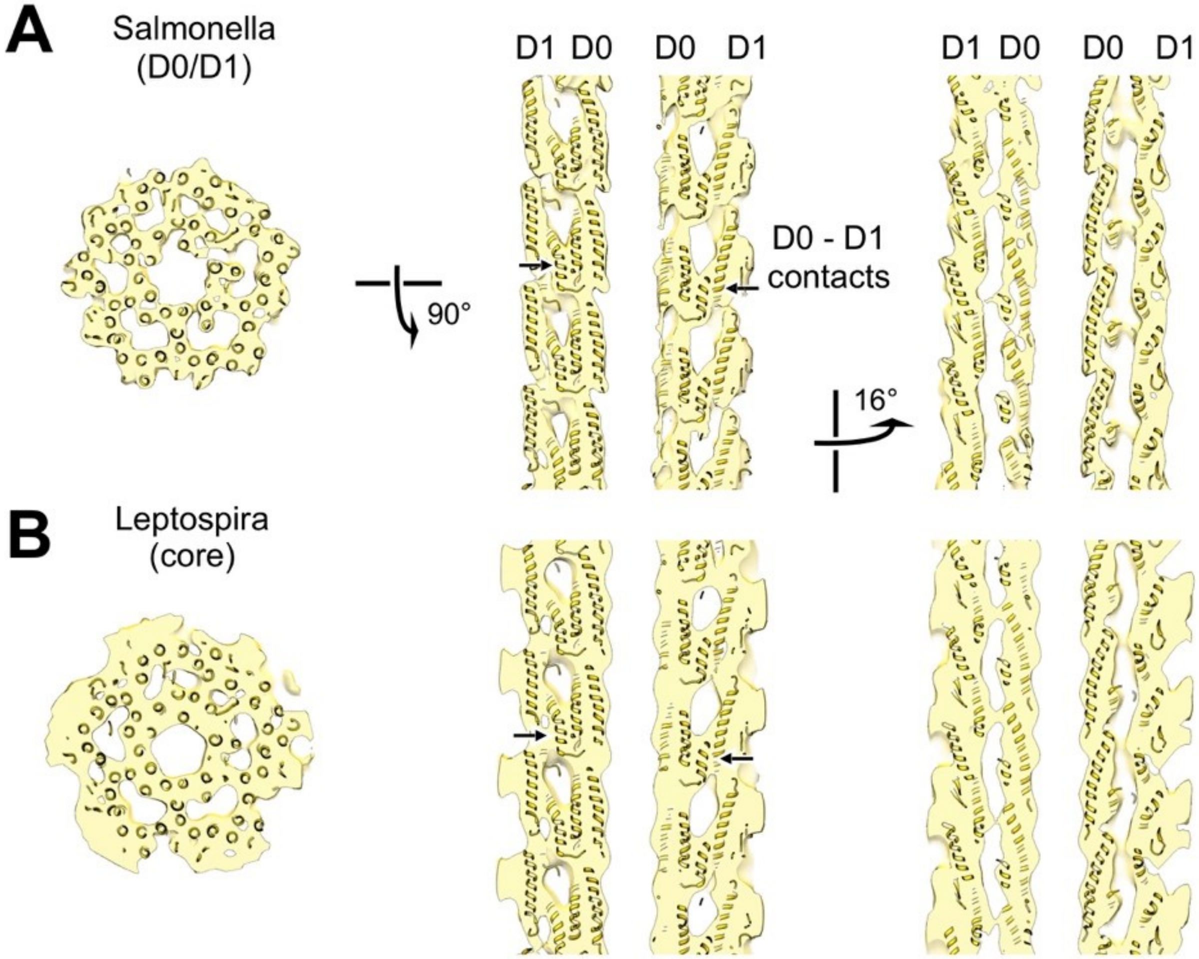


**Supplementary Figure 11**(related to Figure 3). Structural homology between the D0/D1 core region from a synthetic map of the *Salmonella* flagellar filament and the core region of our wild-type *Leptospira* flagellar filament subtomogram average volume. **A.** Axial and lateral cross sections of the *Salmonella* flagellar filament from an atomic model(Yonekura et al., 2003) rendered at 12Å resolution. **B.** Corresponding cross sections of the aligned *Leptospira* flagellar filament subtomogram average volume, after 11-fold symmetry averaging.

**Supplementary Table 1. Protein components of the flagellar filament from *Leptospira*.**

| **Protein** | **Mol. weight** | **Putative localization** in spirochete filaments | **Gene names *Leptospira biflexa*** (Uniprot ID) | **Gene names *Leptospira interrogans*** (Uniprot ID) | **Protein copies per cell in *L. interrogans*** (Malmstrom 2009 *Nature* 460:762) | **Sequence homologies** |
|---|---|---|---|---|---|---|
| **FlaB1** | ~31kDa | Core | LEPBIa2133 (B0SSZ5) | LIC11890 (Q8F4M3) | ~14,000 | • The four FlaB isoforms are homologous to domains D0+D1 of *S. enterica* flagellin FliC.<br>• FlaB1, FlaB2 and FlaB4 are ~65-70% identical to each other in both *Leptospira* species.<br>• *L. biflexa* FlaB1 is ~87% identical to *L. interrogans* FlaB1. |
| **FlaB2** | ~31kDa | Core | LEPBIa2132 (B0SSZ4) | LIC11889 (Q72R59) | ~2,000 | • *L. biflexa* FlaB2 is ~78% identical to *L. interrogans* FlaB2. |
| **FlaB3** | ~31kDa | Core | LEPBIa1872 (B0SS86) | LIC11532 (Q72S54) | ~300 | • FlaB3 is ~50-55% identical to the other three isoforms in both *Leptospira* species.<br>• *L. biflexa* FlaB3 is ~62% identical to *L. interrogans* FlaB3. |
| **FlaB4** | ~31kDa | Core | LEPBIa1589 (B0SQZ5) | LIC11531 (Q72S55) | ~3,500 | • *L. biflexa* FlaB4 is ~92% identical to *L. interrogans* FlaB4. |
| **FlaA1** | ~36kDa | Sheath | LEPBIa2335 (B0SKT4) | LIC10788 (Q72U74) | ~4,500 | • The two FlaA isoforms are not homologous to FlaB or other bacterial flagellins.<br>• FlaA1 and FlaA2 are ~25-28% identical in both *Leptospira* species.<br>• *L. biflexa* FlaA1 is ~56% identical to *L. interrogans* FlaA1. |
| **FlaA2** | ~27kDa | Sheath | LEPBIa2336 (B0SKT5) | LIC10787 (Q72U75) | ~3,500 | • *L. biflexa* FlaA2 is ~70% identical to *L. interrogans* FlaA2. |
| **FcpA** | ~36kDa | Sheath | LEPBIa0267 (B0STJ8) | LIC13166 (Q72MM7) | ~8,000 | • FcpA is unique to the *Leptospira* genus.<br>• FcpA and FcpB are not homologous.<br>• *L. biflexa* FcpA is ~77% identical to |
| **FcpB** | ~32kDa | Sheath | LEPBIa1597 (B0SR03) | LIC11848 (Q72RA0) | ~4,000 | • FcpB is unique to the *Leptospira* genus.<br>• *L. biflexa* FcpB is ~53% identical to *L. interrogans* FcpB. |

**Supplementary Table 2. X-ray diffraction data processing and model refinement statistics.**

| | **FcpA_1** | **FcpA_2** | **FcpA_3** | **FcpB** |
|---|---|---|---|---|
| Wavelength | 0.97910 | 0.97858 | 0.97858 | 1.54179 |
| Resolution range | 67.36 – 1.90<br>(1.94 - 1.90)* | 48.23 – 2.95<br>(3.07 - 2.95) | 45.12 - 2.5<br>(2.6 - 2.5) | 37.15 – 2.58<br>(2.72 - 2.58) |
| Space group | P 622 | $P\ 2_1$ | C 2 | $P\ 2_1 2_1 2_1$ |
| Unit cell (abc Å, αβγ º) | a=b=132.4<br>c=67.4<br><br>α=β=90 γ=120 | a=85.5 b=96.5<br>c=121.1<br><br>α=90 β=105.2 γ=90 | a=82.3 b=99.6<br>c=106.7<br><br>α=90 β=91.9 γ=90 | a=60.7 b=65.6<br>c=134.4<br><br>α=90 β=90 γ=90 |
| Total reflections | 180402 (11851) | 137503 (15749) | 99658 (10749) | 62470 (8683) |
| Unique reflections | 27914 (1742) | 39975 (4485) | 29300 (3278) | 17479 (2442) |
| Multiplicity | 6.5 (6.8) | 3.4 (3.5) | 3.4 (3.3) | 3.6 (3.6) |
| Completeness | 99.7 (99.3) | 99.5 (99.7) | 98.4 (97.9) | 99.2 (97.1) |
| Mean I/sigma(I) | 23.9 (1.5) | 17.2 (2.0) | 18.2 (2.3) | 8.1 (3.2) |
| Wilson B factor | 31.7 | 102.8 | 81.3 | 42.8 |
| R-merge | 0.152 (2.128) | 0.050 (0.651) | 0.040 (0.484) | 0.128 (0.395) |
| R-meas | 0.166 (2.309) | 0.059 (0.768) | 0.047 (0.577) | 0.150 (0.464) |
| $CC_{1/2}$ | 0.991 (0.218) | 0.999 (0.842) | 0.999 (0.805) | 0.993 (0.818) |
| Reflections used in refinement | 27881 (1605) | 39955 (4341) | 29292 (3123) | 17436 (2411) |
| Reflections used for R-free | 1494 (100) | 2005 (208) | 1489 (155) | 1029 (168) |
| R-work | 0.194 (0.259) | 0.197 (0.3445) | 0.196 (0.2586) | 0.204 (0.2739) |
| R-free | 0.219 (0.309) | 0.222 (0.3888) | 0.221 (0.3088) | 0.250 (0.3328) |
| Number of non-hydrogen atoms | 2307 | 7971 | 4089 | 3472 |
| macromolecules | 1979 | 7924 | 3866 | 3410 |
| ligands | 90 | 36 | 84 | 31 |
| solvent | 238 | 11 | 139 | 31 |
| RMS bonds (Å) | 0.010 | 0.010 | 0.010 | 0.010 |
| RMS angles (º) | 0.86 | 1.00 | 1.03 | 1.11 |
| Ramachandran favored ‡ (%) | 98.72 | 98.39 | 98.23 | 95.33 |
| Ramachandran outliers ‡ (%) | 0.00 | 0.00 | 0.22 | 0.00 |
| PDB ID | 6NQW | 6NQX | 6NQY | 6NQZ |
| Raw diffraction data ¶ (doi) | 10.15785/<br>SBGRID/693 | 10.15785/<br>SBGRID/691 | 10.15785/<br>SBGRID/692 | 10.15785/SBGRID/694<br>(data used to solve the structure by SAD)<br><br>10.15785/SBGRID/695<br>(data used for final structure refinement) |

*Values in parentheses are for highest-resolution shell.

‡ Calculated by MolProbity [Williams, C. J. et al. MolProbity: More and better reference data for improved all-atom structure validation. Protein Sci 27, 293-315, doi:10.1002/pro.3330 (2018)]

¶ Deposited in the SBGrid Data Bank public database [Morin, A. et al. Collaboration gets the most out of software. Elife 2, e01456, doi:10.7554/eLife.01456 (2013)].

**Supplementary Table 3. Diameter changes in wild-type and *fcpA*$^{-}$/*fcpB*$^{-}$ mutants reflect differences in sheath composition.**

| *Leptospira* strain | Inner Curvature[1] | Outer Curvature[2] | Both[3] | Total # Images |
|---|---|---|---|---|
| WT | 0 | 4 | 7 | 162 |
| *fcpB*$^{-}$ | 0 | 4 | 2 | 139 |
| *fcpA*$^{-}$ | 16 | 3 | 1 | 83 |

[1] 'Inner curvature' refers to filaments where an abrupt transition in the apparent filament diameter was observed on the concave side of a curved filament.

[2] 'Outer curvature' refers to filaments where an abrupt transition was observed on the convex side.

[3] 'Both' refers to cases where both 'inner' and 'outer' transitions were observed in the same filament (see Supplementary Figure 9).

### SUPPLEMENTARY VIDEOS

**Supplementary Video 1**. Overview of the wild-type *Leptospira* flagellar filament map and model, illustrating separately the FcpA and FcpB sheath layers.

**Supplementary Video 2.** Cross-section of the filament center, showing superposed wild-type map and atomic model, going through a 360° axial rotation of the filament. The core density is replaced by the 11-fold symmetry-averaged map.

**Supplementary Video 3.** Similar to Supplementary Video 2, but without the 11-fold symmetry-averaged core. The core atomic model is not fully accounted for by the map density at certain viewing angles, which we attribute to missing wedge artifacts.

### References:


Agulleiro, J. I., & Fernandez, J. J. (2011). Fast tomographic reconstruction on multicore computers. *Bioinformatics, 27*(4), 582-583. doi:10.1093/bioinformatics/btq692

Beck, T., Krasauskas, A., Gruene, T., & Sheldrick, G. M. (2008). A magic triangle for experimental phasing of macromolecules. *Acta Crystallogr D Biol Crystallogr, 64*(Pt 11), 1179-1182. doi:10.1107/S0907444908030266

Bharat, T. A., Russo, C. J., Lowe, J., Passmore, L. A., & Scheres, S. H. (2015). Advances in Single-Particle Electron Cryomicroscopy Structure Determination applied to Sub-tomogram Averaging. *Structure, 23*(9), 1743-1753. doi:10.1016/j.str.2015.06.026

Bricogne, G., Vonrhein, C., Flensburg, C., Schiltz, M., & Paciorek, W. (2003). Generation, representation and flow of phase information in structure determination: recent developments in and around SHARP 2.0. *Acta Crystallogr D Biol Crystallogr, 59*(Pt 11), 2023-2030. doi:10.1107/s0907444903017694

Bricogne G., B. E., Brandl M., Flensburg C., Keller P., Paciorek W.,, & Roversi P, S. A., Smart O.S., Vonrhein C., Womack T.O. (2017). BUSTER v. 2.10.3. Retrieved from https://www.globalphasing.com/buster/

Cope, J., Heumann, J., & Hoenger, A. (2011). Cryo-electron tomography for structural characterization of macromolecular complexes. *Curr Protoc Protein Sci, Chapter 17*, Unit17 13. doi:10.1002/0471140864.ps1713s65

Crenshaw, H. C., Ciampaglio, C. N., & McHenry, M. (2000). Analysis of the three-dimensional trajectories of organisms: estimates of velocity, curvature and torsion from positional information. *J Exp Biol, 203*(Pt 6), 961-982. Retrieved from https://www.ncbi.nlm.nih.gov/pubmed/10683157

Das, R., & Baker, D. (2008). Macromolecular modeling with rosetta. *Annu Rev Biochem, 77*, 363-382. doi:10.1146/annurev.biochem.77.062906.171838

Emsley, P., Lohkamp, B., Scott, W. G., & Cowtan, K. (2010). Features and development of Coot. *Acta Crystallogr D Biol Crystallogr, 66*(Pt 4), 486-501. doi:10.1107/S0907444910007493

Evans, P. R. (2011). An introduction to data reduction: space-group determination, scaling and intensity statistics. *Acta Crystallogr D Biol Crystallogr, 67*(Pt 4), 282-292. doi:10.1107/S090744491003982X

Himes, B. A., & Zhang, P. (2018). emClarity: software for high-resolution cryo-electron tomography and subtomogram averaging. *Nat Methods, 15*(11), 955-961. doi:10.1038/s41592-018-0167-z

Huehn, A., Cao, W., Elam, W. A., Liu, X., De La Cruz, E. M., & Sindelar, C. V. (2018). The actin filament twist changes abruptly at boundaries between bare and cofilin-decorated segments. *J Biol Chem, 293*(15), 5377-5383. doi:10.1074/jbc.AC118.001843

Kabsch, W. (2010). Xds. *Acta Crystallogr D Biol Crystallogr, 66*(Pt 2), 125-132. doi:S0907444909047337 [pii]

10.1107/S0907444909047337

Kovacs, J. A., Galkin, V. E., & Wriggers, W. (2018). Accurate flexible refinement of atomic models against medium-resolution cryo-EM maps using damped dynamics. *BMC Struct Biol, 18*(1), 12. doi:10.1186/s12900-018-0089-0

Lamzin, V. S., Perrakis, A., & Wilson, K. S. (2012). Crystallography of biological macromolecules. In E. Arnold, D. M. Himmel, & M. G. Rossmann (Eds.), *International Tables for Crystallography* (2nd ed., Vol. Volume F, pp. 525-528): Wiley Online Library.

Mastronarde, D. N. (2005). Automated electron microscope tomography using robust prediction of specimen movements. *J Struct Biol, 152*(1), 36-51. doi:10.1016/j.jsb.2005.07.007

Mastronarde, D. N., & Held, S. R. (2017). Automated tilt series alignment and tomographic reconstruction in IMOD. *J Struct Biol, 197*(2), 102-113. doi:10.1016/j.jsb.2016.07.011

McCoy, A. J., Grosse-Kunstleve, R. W., Adams, P. D., Winn, M. D., Storoni, L. C., & Read, R. J. (2007). Phaser crystallographic software. *J Appl Crystallogr, 40*(Pt 4), 658-674. doi:10.1107/S0021889807021206

Namba, K., Yamashita, I., & Vonderviszt, F. (1989). Structure of the core and central channel of bacterial flagella. *Nature, 342*(6250), 648-654. doi:10.1038/342648a0

Pettersen, E. F., Goddard, T. D., Huang, C. C., Couch, G. S., Greenblatt, D. M., Meng, E. C., & Ferrin, T. E. (2004). UCSF Chimera--a visualization system for exploratory research and analysis. *J Comput Chem, 25*(13), 1605-1612. doi:10.1002/jcc.20084

Rodriguez, D. D., Grosse, C., Himmel, S., Gonzalez, C., de Ilarduya, I. M., Becker, S., . . . Uson, I. (2009). Crystallographic ab initio protein structure solution below atomic resolution. *Nat Methods, 6*(9), 651-653. doi:10.1038/nmeth.1365

San Martin, F., Mechaly, A. E., Larrieux, N., Wunder, E. A., Jr., Ko, A. I., Picardeau, M., . . . Buschiazzo, A. (2017). Crystallization of FcpA from Leptospira, a novel flagellar protein that is essential for pathogenesis. *Acta Crystallogr F Struct Biol Commun, 73*(Pt 3), 123-129. doi:10.1107/S2053230X17002096

Scheres, S. H. (2012). RELION: implementation of a Bayesian approach to cryo-EM structure determination. *J Struct Biol, 180*(3), 519-530. doi:10.1016/j.jsb.2012.09.006

Schneider, T. R., & Sheldrick, G. M. (2002). Substructure solution with SHELXD. *Acta Crystallogr D Biol Crystallogr, 58*(Pt 10 Pt 2), 1772-1779. doi:10.1107/s0907444902011678

Schrodinger, LLC. (2015). *The PyMOL Molecular Graphics System, Version 2.1.0*.

Wang, F., Burrage, A. M., Postel, S., Clark, R. E., Orlova, A., Sundberg, E. J., . . . Egelman, E. H. (2017). A structural model of flagellar filament switching across multiple bacterial species. *Nat Commun, 8*(1), 960. doi:10.1038/s41467-017-01075-5

Winn, M. D., Ballard, C. C., Cowtan, K. D., Dodson, E. J., Emsley, P., Evans, P. R., . . . Wilson, K. S. (2011). Overview of the CCP4 suite and current developments. *Acta Crystallogr D Biol Crystallogr, 67*(Pt 4), 235-242. doi:10.1107/S0907444910045749

Wriggers, W. (2012). Conventions and workflows for using Situs. *Acta Crystallogr D Biol Crystallogr, 68*(Pt 4), 344-351. doi:10.1107/S0907444911049791

Wunder, E. A., Jr., Figueira, C. P., Benaroudj, N., Hu, B., Tong, B. A., Trajtenberg, F., . . . Ko, A. I. (2016). A novel flagellar sheath protein, FcpA, determines filament coiling, translational motility and virulence for the Leptospira spirochete. *Mol Microbiol, 101*(3), 457-470. doi:10.1111/mmi.13403

Wunder, E. A., Jr., Slamti, L., Suwondo, D. N., Gibson, K. H., Shang, Z., Sindelar, C. V., . . . Picardeau, M. (2018). FcpB Is a Surface Filament Protein of the Endoflagellum Required for the Motility of the Spirochete Leptospira. *Front Cell Infect Microbiol, 8*, 130. doi:10.3389/fcimb.2018.00130

Yonekura, K., Maki-Yonekura, S., & Namba, K. (2003). Complete atomic model of the bacterial flagellar filament by electron cryomicroscopy. *Nature, 424*(6949), 643-650. doi:10.1038/nature01830
